# Synchronized cardiac impulses emerge from multi-scale, heterogeneous local calcium signals within and among cells of heart pacemaker tissue

**DOI:** 10.1101/2020.04.14.039461

**Authors:** Rostislav Bychkov, Magdalena Juhaszova, Kenta Tsutsui, Christopher Coletta, Michael D. Stern, Victor A. Maltsev, Edward G. Lakatta

**Author notes:** **Contact information:** Edward G. Lakatta, M.D., Laboratory of Cardiovascular Science, NIA/NIH, Biomedical Research Center, 251 Bayview Blvd., Baltimore, MD 21224, USA.

## Abstract

**Background:** The current paradigm of Sinoatrial Node (SAN) impulse generation: (i) is that full-scale action potentials (APs) of a common frequency are initiated at one site and are conducted within the SAN along smooth isochrones; and (ii) does not feature fine details of Ca^2+^ signalling present in isolated SAN cells, in which small subcellular, subthreshold local Ca^2+^ releases (LCRs) self-organize to generate cell-wide APs.

**Objectives:** To study subcellular Ca^2+^ signals within and among cells comprising the SAN tissue.

**Methods:** We combined immunolabeling with a novel technique to detect the occurrence of LCRs and AP-induced Ca^2+^ transients (APCTs) in individual pixels (chonopix) across the entire mouse SAN images.

**Results:** At high magnification, Ca^2+^ signals appeared markedly heterogeneous in space, amplitude, frequency, and phase among cells comprising an HCN4^+^/CX43^-^ cell meshwork. The signalling exhibited several distinguishable patterns of LCR/APCT interactions within and among cells. Apparently conducting rhythmic APCTs of the meshwork were transferred to a truly conducting HCN4^-^/CX43^+^ network of straited cells via narrow functional interfaces where different cell types intertwine, i.e. the SAN anatomical/functional unit. At low magnification, the earliest APCT of each cycle occurred within a small area of the HCN4 meshwork and subsequent APCT appearance throughout SAN pixels was discontinuous.

**Conclusions:** We have discovered a novel, microscopic Ca^2+^ signalling paradigm of SAN operation that has escaped detection using low-resolution, macroscopic tissue isochrones employed in prior studies: APs emerge from heterogeneous subcellular subthreshold Ca^2+^ signals, resembling multiscale complex processes of impulse generation within clusters of neurons in neuronal networks.

**Condensed abstract:** By combining immunolabeling with a novel optical technique we detected markedly heterogenous Ca^2+^signals within and among cell clusters of an HCN4^+^/CX43^-^ meshwork in mouse sinoatrial node. These Ca^2+^ signals self-organized and transferred, throughout the node, to projections from an HCN4^-^/CX43^+^ network connected to a highly organized, rapidly conducting part of the CX43^+^ network. Thus, APs emerge from heterogeneous, subthreshold Ca^2+^ signaling not detected in low-resolution macroscopic isochrones. Our discovery requires a fundamental paradigm shift from concentric impulse propagation initiated within a leading site, to a multiscale/complex process, resembling the emergence of organized signals from heterogeneous local signals within neuronal networks.

## Introduction

The sinoatrial node (SAN), the heart’s primary pacemaker, exhibits a high degree of structural and functional complexity that is vital for generating flexible heart rates and robust rhythms (1). Following the discovery of the SAN in 1906 by Keith and Flack (2) numerous SAN cell characteristics have been described including a diversity of: cell shapes and protein expression; arrangement of cells within SAN tissue (e.g. gradient vs. mosaic (3)); autonomic neuronal input (4); cell to cell electrical and mechanical interactions (5); and intranodal impulse initiation, spread within, and exit from, the SAN (6–9).

A generalized view of coordinated firing of pacemaker cells within the SAN emerged about 40 years ago, in which a dominant or “master” pacemaker cell or a leading pacemaker center dictates the excitation rate and rhythm of thousands of other SAN pacemaker cells by overdriving their intrinsic spontaneous excitation rates (6,10). Shortly thereafter, however, an idea of mutual entrainment of coupled oscillators (11) was applied to the coordinated firing of the entire population of SAN cells (12,13): individual SAN cells that are loosely connected through low resistance junctions generate spontaneous excitations that differ in phase, mutually entrain each other to fire with a common period. The respective intrinsic cell oscillation frequencies of individual cells and the degree of intercellular electrical coupling determined the common period at which all cells fire. In other terms, the mutual entrainment theory of SAN (12) proposed that while the ensemble of SAN cells can be entrained to operate at a given frequency, there can be marked phase differences among spontaneous excitations of individual cells, and the frequency of impulses that exit the SAN lies somewhere between the fastest and slowest spontaneous intrinsic excitation rates of resident SAN cells.

Thus, rather mimicking classical electrical conduction by consecutively exciting each other, as in ventricular muscle tissue, SAN cells are in fact mutually entrained by phase resetting and conduction within SAN is only “apparent” (13,14). The idea of apparent conduction was a major departure from the classic view of the SAN as a slowly conducting tissue in which the action potential (AP), initiated by a small cluster of dominant pacemaker cells, propagates radially at an accelerating pace. This idea was supported by later studies that showed that the initiation of the cardiac impulse could be multimeric, i.e. initiated from different locations (15). The early ideas of mutual entrainment were further elaborated in elegant studies by Verheijck et al. (16) using computer-controlled coupling conductance between individual pacemaker cells and found a critical coupling conductance for 1:1 frequency entrainment of less than 0.5 nS (i.e. generated by a few connexin molecules).

Similar to the evolution of ideas on how the SAN impulse is generated, the functional paradigm of pacemaker cells operating in isolation has also undergone substantial refinement from a simple to more complex origin: from what was described as **the** “pacemaker channel” that solely drives spontaneous diastolic depolarization (17), to a modern concept of a coupled-system of a chemical and ion current oscillators (18,19). According to this coupled oscillator theory, rhythmic local intracellular Ca^2+^ releases (LCRs) generated by an intracellular Ca^2+^ oscillator during diastole self-organize into a powerful and timely Ca^2+^ signal that drives the ensemble of membrane current oscillators to generate inward current culminating in membrane depolarization, sufficient to trigger a cell-wide AP.

More specifically, the interactions of spontaneous LCRs with Na^+^/Ca^2+^ exchanger (NCX) begin to occur near the maximum diastolic potential in the context of NCX current activation and anomalous rectification of HCN4 current both activated by hyperpolarization, and decay of K^+^ currents. These combined actions of LCRs, NCX, and HCN4 regulate cell surface membrane potential within a range that activates low-threshold I_CaL_ (Cav1.3), or I_CaT_ (Cav3.1) which, in turn, via Ca^2+^-influx-induced Ca^2+^ release via ryanodine receptors, increases and feeds forward the ensemble LCR Ca^2+^ signal and NCX current to accelerate the rate of the diastolic depolarization (20). The transition in LCR characteristics is steeply nonlinear, resembling a phase transition (21). Thus, simultaneous growth of the diastolic ensemble LCR signal and diastolic surface membrane depolarization are thus manifestations of a slowly evolving self-organized electrochemical gradient oscillation that reaches criticality in late diastole, followed by a phase transition manifest as the rapid AP upstroke that triggers a whole cell cytosolic Ca^2+^ transient, i.e. AP-induced Ca^2+^ transient (APCT). In other terms, spontaneous LCR emergence at a critical time of an AP cycle (late diastole) entrains membrane depolarization, and late diastole, therefore, might be considered to be an “intrinsic entrainment zone” (14) or a “time window” in which chemical and current oscillators become mutually entrained, controlling the rate and rhythm of AP firing of that cell.

A wide spectrum of spontaneous AP firing cycle lengths observed among single SAN cells in isolation under a large variety of experimental conditions is, in fact, predicted by the spectrum and degree of synchronization of LCR periods among these cells (19,22). Here we hypothesized that on the larger scale of intact SAN tissue LCRs not only occur **within** cells, but (i) differ in spatial distribution, frequency, amplitude, and phase; and (ii) that heterogeneity of local Ca^2+^ signalling characteristics **among** cells translates into different pattern of pacemaker cell excitation within the node.

## Methods

Our experiments conformed to the Guide for the Care and Use of Laboratory Animals, published by the US National Institutes of Health. The experimental protocols were approved by the Animal Care and Use Committee of the National Institutes of Health (protocol #034-LCS-2019). We used one-to-three month old C57BL mice (Charles River Laboratories, USA) anesthetized with sodium pentobarbital (50 mg/kg). The adequacy of anesthesia was monitored until reflexes to tail pinch were lost.

### Sinoatrial Node Preparation

The SAN was dissected according to standard methods (23). The heart was removed quickly and placed in standard Tyrode solution containing (in mM): 130 NaCl, 24 NaHCO_3_, 1.2 NaH_2_PO_4_, 1.0 MgCl_2_, 1.8 CaCl_2_, 4.0 KCl, 5.6 glucose equilibrated with 95% O_2_ / 5% CO_2_ (pH 7.4 at 35.5°C). The whole heart was pinned to a silicon platform under a surgical microscope in order to excise the right and left atria. A 10-ml tissue bath was perfused with standard solution at a rate of 10 ml/min. After removal of the ventricles, the right atrium was opened to expose the crista terminalis, the inter-caval area, and the inter-atrial septum. The preparation was not trimmed, leaving SAN region together with surrounding atria and superior and inferior vena cava (SVC and IVC) intact. The SAN preparation was pinned to the silicon bottom of the experimental chamber by small stainless-steel pins with the endocardial side exposed. Care was taken to provide the minimal amount of stretch required to flatten the SAN tissue. After mounting, the preparation was superfused with solution maintained at a temperature of 36±0.3°C. Anatomic landmarks were used to locate the SAN (Fig 1A).

**Fig 1.**
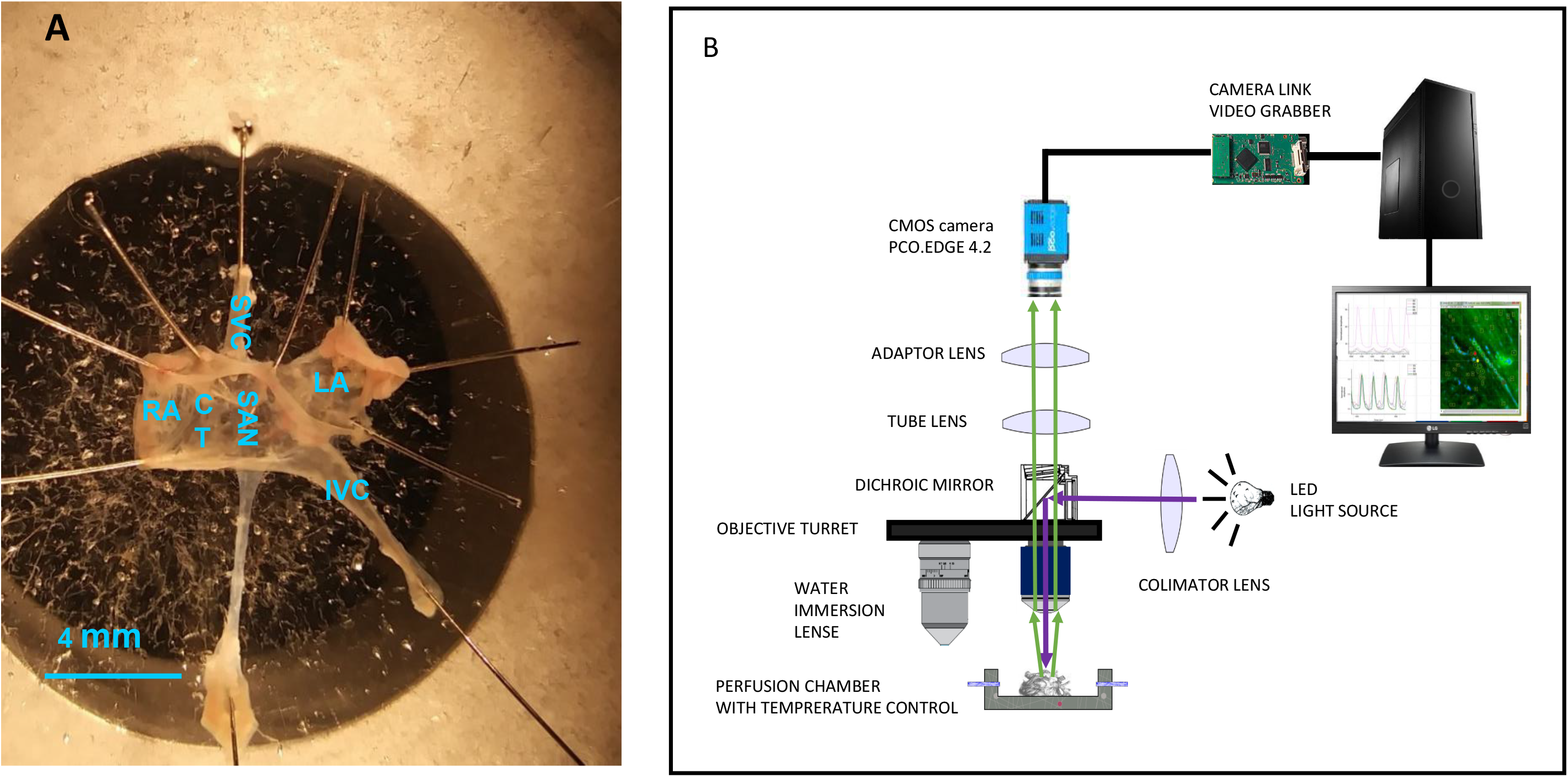
**Panel A:** A SAN preparation pinned to the sylgard silicone bottom of a petri dish fixed to a microscope table. SVC-superior vena cava, IVC inferior vena cava, CT-crista terminalis, RA-right atrium, LA-left atrium. **Panel B:** A schematic representation of the experimental setup for macroscopic and microscopic optical mapping. See methods for details.

### Optical system for SAN imaging

We developed a novel imaging system to assess intracellular Ca^2+^ dynamics within individual cells resident within the entire mouse intact SAN. We used a stationary fixed stage upright microscope (AxioExaminer D1 equipped with zoom tube (0.5–4x), Carl Zeiss Microscopy LLC) and a camera (sCMOS PCO edge 4.2) with a high spatial and temporal resolution (Fig 1B). The microscope was mounted on a motorized X-Y MT-2078/MT-2278 translator (Sutter Instruments). The experimental chamber with the SAN preparation was placed on a platform (Sutter Instruments) that was mounted onto a pressurized air table (Newport).

### Imaging of local Ca^2+^ signals in SAN tissue

The SAN preparation was incubated with a membrane-permeable Ca^2+^ indicator Fluo-4AM (10μM) for 1.5 hours. In some experiments we also recorded Ca^2+^ signals specific to HCN4^+^ cells using SANs from mice (pCAGGS-GCaMP8) in which Ca^2+^ signals were driven by HCN4-promoter, provided by Michael Kotlikoff VMD, PhD, College of Veterinary Medicine, Cornell University (24).

The excitation light (CoolLED pE-300ultra) was directed to the microscope via a single port epifluorescence condenser, providing uniform illumination of the object plane. To optimize the signal to noise ratio and durability of the preparation, we applied excitation light to the SAN at 40% of the maximum power of the light source CoolLED. The excitation light was reflected to the SAN preparation by a dichroic mirror with a central wavelength of 498 nm, and the emitted fluorescence signal was collected through a 530±20 nm filter (Semrock, USA). The fluorescence image of the SAN preparation was projected by air or water lenses onto the sCMOS camera sensor. To prevent interference of tissue motion during recordings from SAN tissue, we decoupled electrical excitation and mechanical contraction in some preparations by inhibiting the formation of the Ca^2+^-sensitive regulatory complexes within sarcomeres using 10 μM cytochalasin B (25). Otherwise residual mechanical artefacts produced during diastolic phase were compensated with ImageJ software during the post processing of recorded images (https://imagej.nih.gov).

To image APCTs across entire SAN preparations (n=7) from SVC to IVC, we used 2.5x and 5x magnification. To visualize local Ca^2+^ dynamics of individual cells within the SAN in a given Region of Interest (ROI), we used 10x and 20x magnification water-immersion lenses (W Plan-Apochromat 10x/0.5 M27 75mm, Plan-Apochromat 20x/1.0 DIC D=0.17 M27 75mm). Ca^2+^ signals had a high signal to noise ratio (>10) over the entire SAN from SVC to IVC, even at a low optical resolution with a 2.5x (air) lens. This signal consistency, together with high temporal (500-700 Hz) and spatial resolution of SAN images (2200×320 pixels of 2×4 mm tissue) allowed acquisition of data required to construct spatial and temporal maps of SAN cell network activation in each pixel, i.e. chrono-pixel or “chronopix” for short. Ca^2+^ indicator fluorescence signal intensity was stable for 20-35 minutes.

### Simultaneous Ca^2+^ imaging and AP recording

We used spontaneous APCTs to inform on the occurrence and locations of spontaneous APs within the SAN. To validate APCTs as reporters of APs, we simultaneously measured APs and APCTs in a subset (n=3) of SAN preparations. To record the transmembrane potential, we used sharp microelectrodes (40-70 MΩ) fabricated from aluminosilicate glass capillaries (1.5mm OD 0.86mm ID) with a horizontal pipette puller (Sutter Instruments, CA, USA), and back-filled with 3M KCl. The neck of the recording sharp microelectrode was pulled long enough to be sufficiently “springy” in order to follow tissue movement without compromising the recording. A sharp glass microelectrode was inserted from the endocardial side of the tissue. APs were recorded with a high impedance amplifier (A-M Systems, USA) with a virtual bridge circuit for current injection. The electrical signal was digitized with Digidata 1440 and analyzed with PCLAMP10 software (both from Molecular Dynamics, USA).

### Data analysis

Microsoft Excel and OriginLab software were used to analyze data sets of membrane potential recordings, amplitudes of intracellular Ca^2+^ transients measured within the ROIs, delay times, to normalize to maximum of recorded traces and to make linear fit. Then we also used OriginLab to generate power spectra in time series of local Ca^2+^ signals. The time series were generated from average signal within each ROI for each frame in the sequentially recorded stack of images. Note that while we present our data in terms of chronopix depicting the time in ms at which Ca signal follow the occurrence of the earliest Ca signal, they can be also expressed in radians or percentage of the cycle length as stated in the coupled oscillator theory.

### Whole mount SAN Immunolabeling

We combined Ca^2+^ imaging with immunolabeling to correlate Ca^2+^ dynamics with cytoarchitechture. Another subset (n=7) of SAN preparations was fixed in 4% paraformaldehyde overnight at 4°C. The SANs were washed three times in phosphate-buffered saline (PBS) and permeabilized overnight in PBS containing 0.2% Triton X-100 and 20% DMSO. After blocking the non-specific binding sites by incubation for 8 hours with 0.2% Tween-20 in PBS containing 3% normal donkey serum, SAN whole-mount preparations were incubated for 3 days with the primary antibodies diluted in PBS containing 0.2% Tween-20 and 3% normal donkey serum. They were then washed three times with PBS containing 0.2% Tween-20, incubated overnight with appropriate secondary antibodies then washed three times with PBS containing 0.2% Tween-20. Whole-mount SAN preparations were mounted in Vectashield (Vector Laboratories) and sealed with a coverslip. The SAN preparations were mounted with the endocardium uppermost. Immunolabeling of whole-mount SANs was imaged with a ZEISS-LSM510 confocal microscope and a ZEISS-AxioExaminer D1 fluorescence microscope with zoom tube (0.5–4x) equipped with appropriate filters for fluorescence spectra. Parts of the whole-mount SAN were imaged individually, and these images were concatenated into one output image to portray entire SAN preparation. To obtain high resolution images of the entire SAN, the preparations were imaged in a tile scanning mode. Fluorescence of immunolabelled cells within SAN tissue in 3D was visualized by confocal optical slicing (Z stacking) to a depth up to 100 μM from the endothelial surface.

Antibodies: HCN4^+^ cells were identified by rabbit polyclonal antibodies for hyperpolarization-activated, cyclic nucleotide-gate cation channels HCN4 (1:250; Alomone Labs). Mouse monoclonal antibody to connexin 43 (1:250; Invitrogen) was used to label the gap junctions of striated cells and Alexa Fluor 633 phalloidin was used to visualize the F-actin filaments.

### Immunolabeling of whole mount SAN for HCN4, connexin 43 and labelling of F-actin

Previous studies in thin slices from central SAN demonstrated that numerous cells have strong HCN4 immunolabeling and lacked CX43 (26). To explore the expression patterns of key molecules/markers of pacemaker cells, we immunolabeled and reconstructed nearly the entire cellular network within intact whole mount SAN preparations. We analyzed immunolabeling of the entire SAN at low optical magnification and analyzed immunolabeling within regions in which leading cells with rhythmical LCRs preceded APCTs at higher magnification. We employed confocal imaging to acquire a 3D reconstruction of the optically dissected intact SAN tissue (of note, in previous studies 3D reconstruction was performed by physical dissection of SAN tissue with a microtome into separate tissue slices). The resultant 3D-networks of intact cells expressing HCN4, F-actin and/or CX43 were reconstructed and further examined with Zeiss imaging software ZEN2. Images were processed with Image J software (https://imagej.nih.gov open-source, NIH)

## Results

### Simultaneous action potential recording and intracellular Ca^2+^ imaging

We imaged spontaneous APCTs to report the occurrence of spontaneous APs. In order to ensure the fidelity between the occurrence of APCTs and the occurrence of APs, we simultaneously acquired Ca^2+^ signal fluorescence, recorded at lower optical power, and APs, recorded with conventional sharp microelectrodes in a subset of experiments (Fig 2). In each experiment, a ROI within a Ca^2+^ image at low magnification was selected around the tip of the sharp microelectrode. Figure 2A shows a representative example of simultaneously recorded AP-s and APCTs. The AP trace and Ca^2+^ transient recordings, normalized to their peak amplitudes, are superimposed in Figure 2B. Note that, as expected, an APCT shortly followed the corresponding AP onset. APCT cycle lengths plotted against AP cycle lengths showed a tight linear relationship (Fig 2C), validating the use of whole mount APCT images to report the occurrence of APs within SAN tissue.

**Fig 2.**
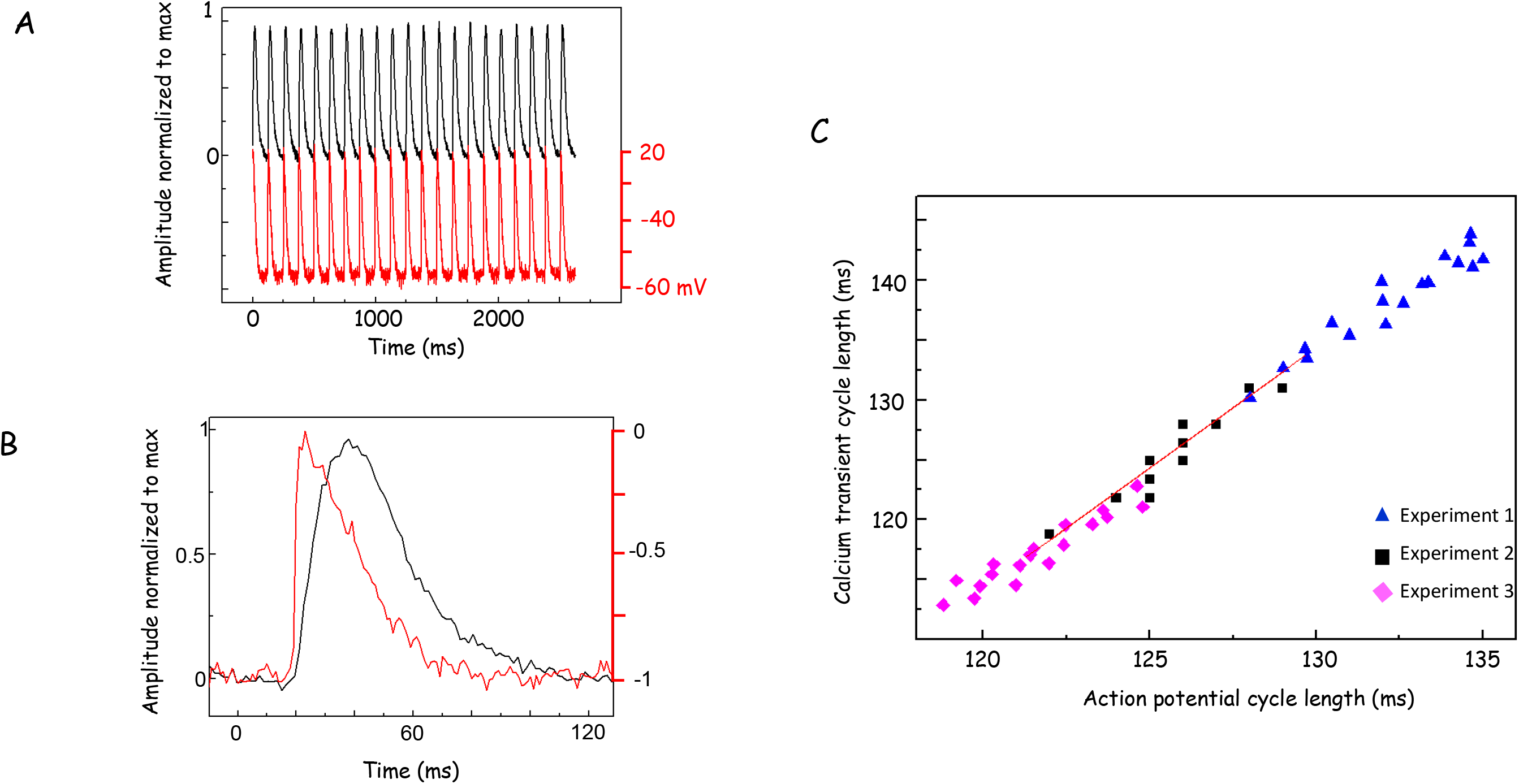
**Panel A:** Continuous simultaneous recordings of action potentials with sharp microelectrode (AP, red color) and action potential-induced Ca^2+^ transients (AP, black color) recorded as Fluo 4 fluorescence transients. APCT fluorescence transients are normalized to maximum. Ca^2+^ and membrane potential were recorded near crista terminalis in the SAN area (not in the auricle site). **Panel B:** Superimposed normalized AP and APCT on an expanded time scale. **Panel C:** APCT cycle lengths faithfully report AP cycle lengths, r^2^=0.9.

### Panoramic Imaging (at 2.5x) of the entire SAN

The mouse SAN averages 100-200 μm in thickness and up to 1000 μm in width and 4000 μm in length, enabling visualization of the entire SAN preparation within a microscopic field of view at low optical magnification. We quantified Ca^2+^ signals in defined ROIs within a 2D SAN image.

Figure 3 illustrates a complete spatiotemporal pattern (red color) of the earliest detectable APCTs and of subsequent APCT occurrences within chronopix across the entire SAN within a given focal plane (Video 1). Two ROIs in panel A that are separated by approximately 300 μm, show the location in which the earliest APCTs emerge (black ROI), and the region where the next APCTs emerged (blue ROI). When the APCTs from the two ROIs (in panel A) were normalized to their maxima, superimposed, and plotted as function of time, their phase difference was 7 ms (Fig 3C). The earliest APCTs occurred within a region of about 20 by 200 μm (black ROI in panel A), near the SVC. This can be clearly seen as a small red spot at the 2 ms chronopix and subsequent APCTs emerged 4-18 ms thereafter (red color in a sequence of chronopix in Fig 3B).

**Fig 3.**
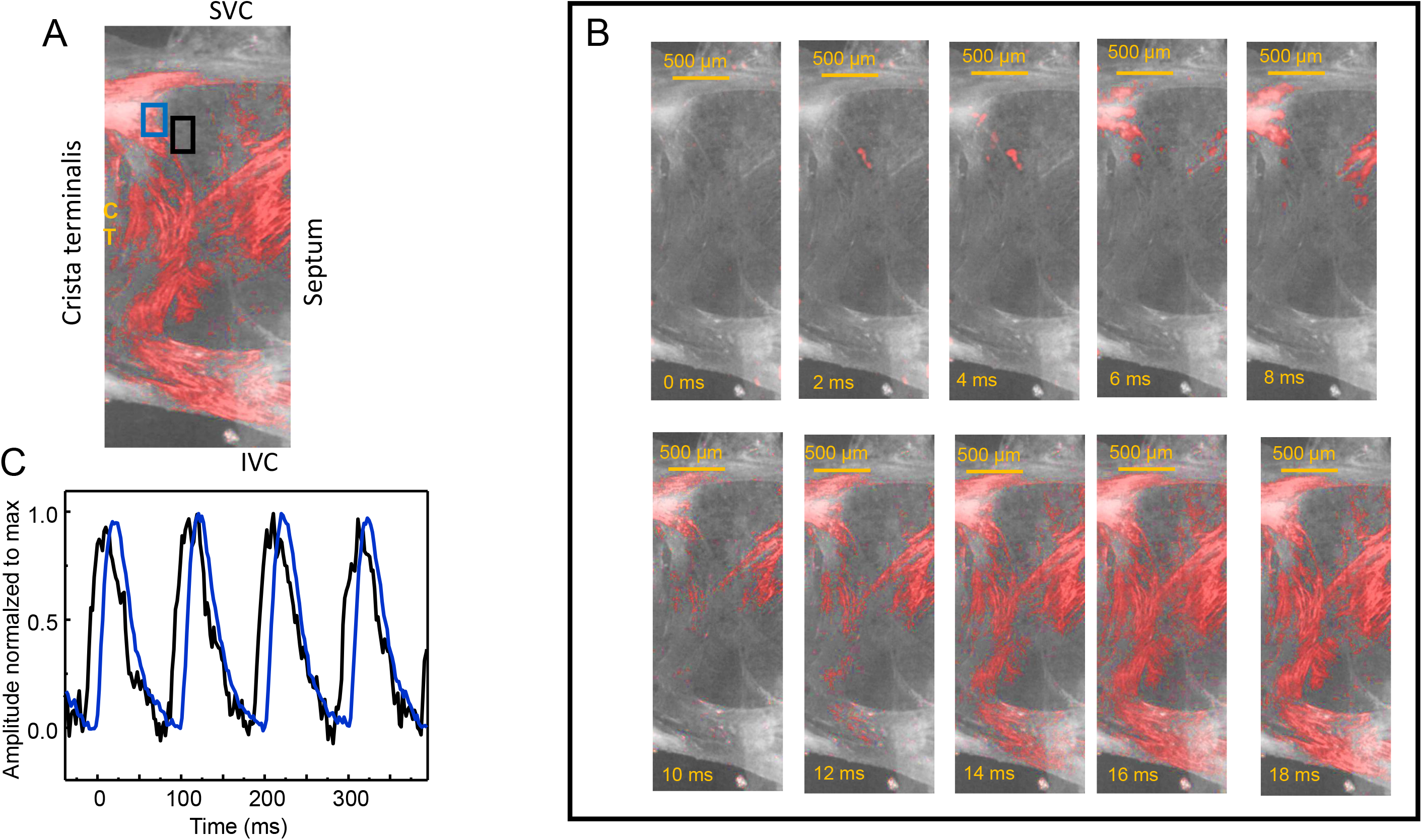
Images showing the sequence of APCTs appearance (red color) within chronopix across the SAN. **Panel A:** A SAN preparation viewed at low optical magnification: SVC-superior vena cava, IVC-inferior vena cava. Two regions of interest (ROIs), identified by phase shift analysis of chronopix data recorded in panel A, show the location of pixels in which the earliest APCTs emerge (black ROI) and chronopix in which the next APCTs emerged (blue ROI). **Panel B:** Series of images acquired every 2 ms depicting the discontinuous emergence of APCTs within the chronopix across the node when viewed in a given focal plane. **Panel C:** Superimposed traces of APCTs obtained in the ROIs of the respective colors (black and blue) in Panel A.

### Heterogeneous LCR-APCT patterns among clusters of SAN cells localized within the central SAN

We further inspected the area where the earliest APCT occurred (blue rectangle in Fig 4A, near to the black ROI in Fig 3) at a higher optical resolution using 10x and 20x water-immersion lenses. We discovered Ca^2+^ signals that were markedly heterogeneous within and among cells ranging from highly synchronized to less synchronized APCTs, manifest as variably synchronized increases in intracellular Ca^2+^ throughout the entire volume of a cell (Fig 4B yellow rectangles). LCRs, in contrast, were manifest as an increase in Ca^2+^ signal in one part of the cell, while at the same time, Ca^2+^ in other parts of the same cell remained at its basal level (Fig 4B red rectangle) (Video 2). Importantly, many cells manifested only low amplitude LCRs, but not APCTs. Also, of note, the earliest Ca^2+^ signal within the ROI did not originate from the same cell in every cycle (not shown).

**Fig 4.**
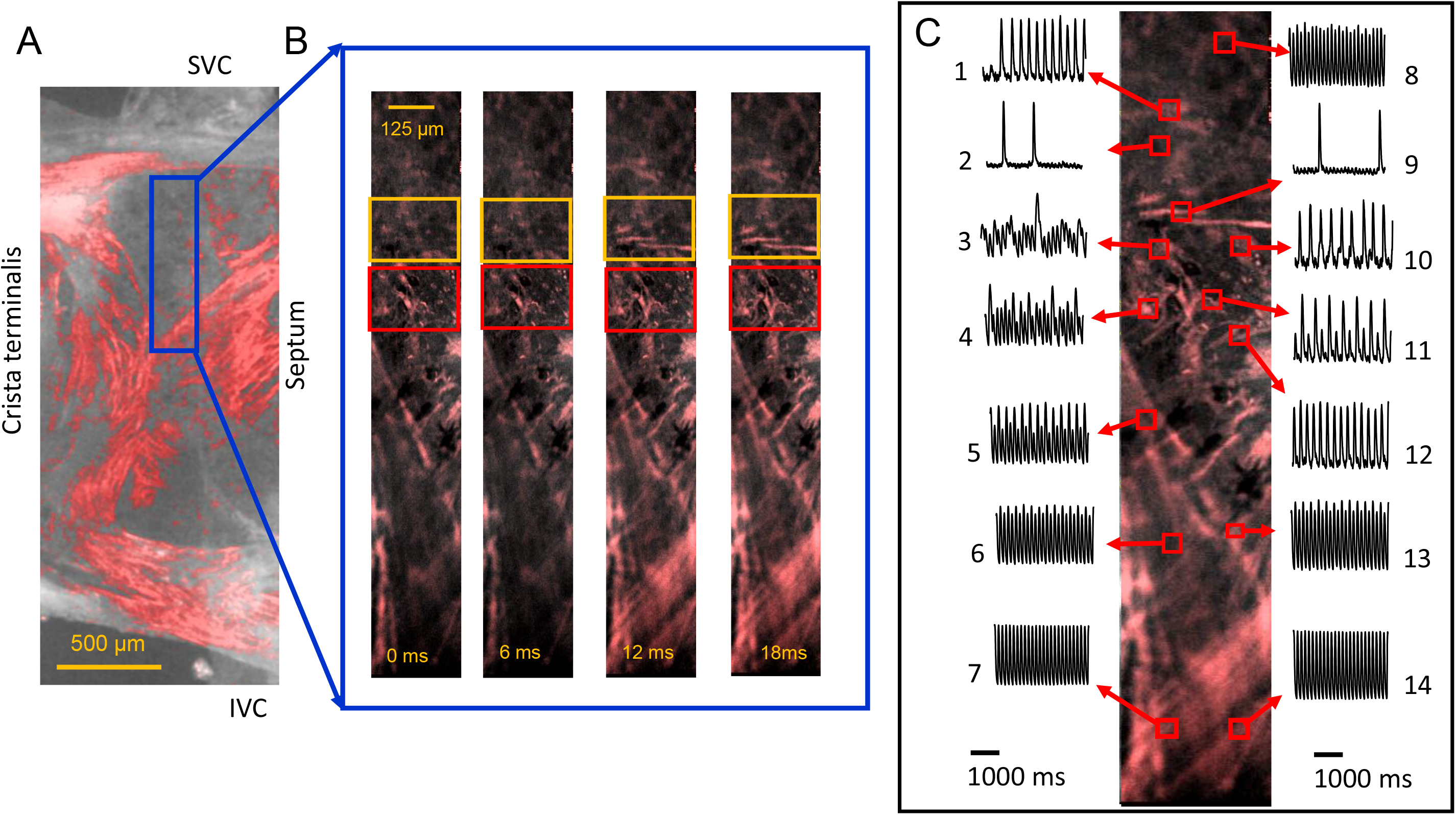
Substantial heterogeneity of local Ca^2+^ signals within the area of the earliest APCT occurrence. **Panel A:** A SAN preparation at low optical magnification: SVC-superior vena cava; IVC inferior vena cava. Blue rectangle shows the location of the ROI within the central part of the SAN scanned at low (10x water immersion objective) optical magnification. **Panel B:** Sequences of Fluo-4 fluorescence images acquired every 6 ms within the blue box in panel A. Yellow rectangles outline SAN cells that generated APCTs and red rectangles outline cells with dominant LCRs but without apparent APCTs. **Panel C:** Heterogeneity of Ca^2+^ signals among cells within the blue rectangle in panels A and B. Red squares indicate ROIs, and arrows point to the respective time series of the average Ca^2+^ signal recorded from cells within those ROIs. The Ca^2+^ signals of different amplitudes within each ROI were normalized to their maximum.

Figure 4, Panel C contrasts the large variety of Ca^2+^ signals generated in cells within the area in which the earliest APCT appears (blue rectangle in panel A). SAN cells in ROIs 1, 2, 8, 9, 10, 11 and 12 in panel C generated **both** APCTs and LCRs; cells in ROIs 6, 7, 13, 14 generated **only** APCTs, but **not** LCRs; while cells in the ROIs 3 and 4 persistently produced LCRs, but **not** APCT’s.

Furthermore, spontaneous intrinsic frequencies and amplitudes of Ca^2+^ signals within different ROIs were markedly heterogeneous:

- Cells within ROIs 7, 8 and 14 generated only APCTs of relatively constant amplitude at a frequency of 8.1Hz, which was also the frequency of AP’s recorded in the right atrium (not shown).
- Cells within ROIs 5, 6 and 13 generated APCTs at frequencies that were also close to those in ROIs 7, 8 and 14, but exhibited APCT amplitude alternans.
- Cells in ROIs 2, 9 and 10 generated APCTs at low frequencies of 0.9 Hz, 0.6 Hz and 2.8Hz), much lower than that in ROI’s 6, 13, 7, 8, 14
- Cells within ROI 11 generated APCTs at a low frequency of 4.7Hz that alternated in amplitude.
- And importantly, the cell in ROI 1 was initially silent, but then began to generate spontaneous APCTs at a frequency 3.7 Hz during the course of sequential image recording. Thus, cells can generate APCTs both steadily and in bursts.

Furthermore, we also detected similar multimodal Ca^2+^ signals of variable amplitudes and durations in SANs from genetically manipulated mice (pCAGGS-GCaMP8) in which Ca signals were driven by HCN4 promoter, i.e. Ca signals specific to HCN4^+^ cells (Video 3).

### LCRs precede APCTs in some cells within the central SAN

To uncover patterns of the relationships between LCRs and APCTs in cells, we focused on diastolic phases of APCT cycles. In some cells we observed a temporal relationship between LCRs and APCTs that is characteristic of single SAN cells in isolation (27), i.e. diastolic LCRs in a given cell occurred prior to the APCT firing in that cell. A typical example is shown in Fig 5 and Video 4. LCRs and APCTs measured within chonopix in the green rectangle (Panel A) are observed as small bright spots occurring during the diastolic phase (yellow arrows in Fig 5B). Line scan images provide a convenient, instant view of Ca^2+^ dynamics (i.e. Ca^2+^ signal vs. time) localized along a selected line within the movie, i.e. in the corresponding sequence of TIF files. The sequence of line-scans (Fig 5C) clearly illustrates LCRs occurrence prior to APCTs in the same cell during several cycles. Overlaid plots of LCRs and APCTs show that LCRs onset occurs during an intrinsic, diastolic “entrainment zone” (14) prior to the onset of APCTs within each cycle (Fig 5D).

**Fig 5.**
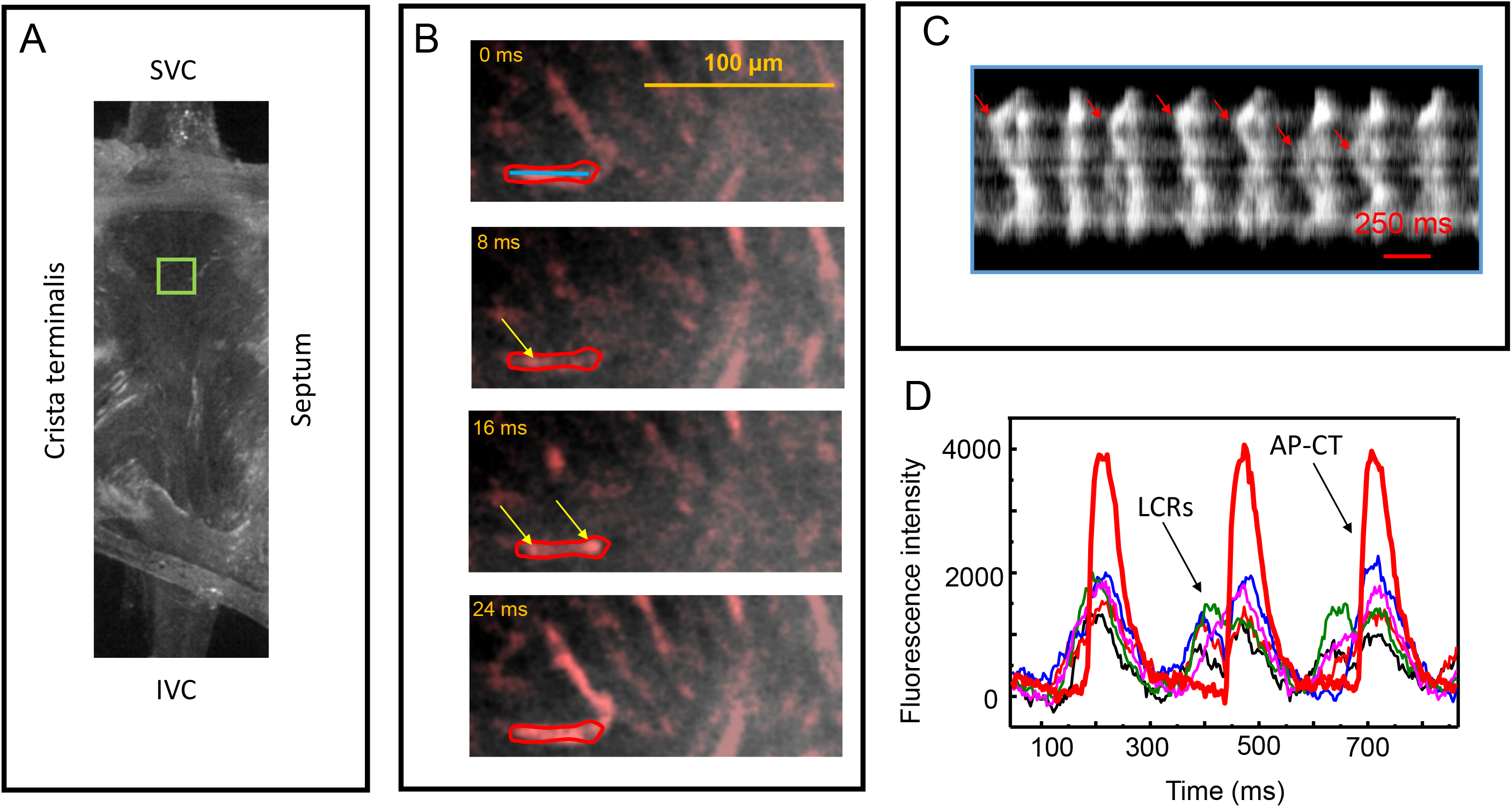
Temporal relationships between LCRs and APCTs generated within the same cell. **Panel A:** A SAN preparation at low optical magnification. **Panel B:** Ca^2+^ signals in SAN cells within the green rectangle in panel A, recorded via a 20x water immersion objective. Spontaneous LCRs, indicated by yellow arrows, appear as bright spots in the cell within the red ROI. **Panel C**. Line scan image of Ca^2+^ signals within the ROI in panel B. Blue line in the top panel B indicates the placement of the line scan. Each cycle exhibits spontaneous LCRs occurring during the diastolic phase. Red arrows on the line scan image point to examples of LCRs. **Panel D:** APCTs (red color) and LCRs (traces with variable colors and smaller amplitudes) in the cell in panels B and C, plotted as function of time.

### Some SAN cells generate only LCRs that precede APCTs in adjacent cells

We observed a novel type of LCR-APCT relationship (Fig 6A, green ROI) in which some cells within a cell cluster did not generate APCTs, but generated only LCRs (Fig 6B, yellow ROI) that preceded APCT firing in one or several adjacent cells that did not manifest their own LCRs (Fig 6B red ROI, Video 5). Note that the cell with only LCRs generated the earliest signal within the cluster. In this cell, having LCRs only, the LCRs emerged during the diastolic period at 16 ms that preceded AP firing of the **<u>adjacent</u>** SAN cells at 24 ms (red ROI Fig 6B). Note also that the cells adjacent to those cells generating only APCTs did not manifest intrinsic LCRs prior to generating a synchronous APCT (Video 5). A sequence of line-scans recorded from the leading cell firing only LCRs and from an adjacent cell that fired APCTs but without LCRs (Fig 6C) confirmed that LCRs in the cell without APCT’s (image within lower yellow frame Fig 6C) occurred prior to APCTs in the adjacent cell (image within top red frame Fig 6C). Superimposed time series of the chronopixs representing LCRs from the cell with LCRs only, and APCTs from an adjacent cell (Fig 6D) also show that LCRs onset in the cell with only LCRs occurred during diastolic phase of adjacent cells. Note that the amplitude of the ensemble LCR Ca^2+^ signal continued to organize until the APCT in the adjacent cell reached its peak amplitude, after which the LCR ensemble Ca^2+^ signal decayed to basal level, indicating that the LCRs Ca^2+^ dynamics in the leading cell, in this case, are not externally reset by AP occurrence in the adjacent neighboring cells. This temporal relationship was observed during all recorded cycles in this preparation and similar behavior was observed in 7 of 7 SAN preparations.

**Fig 6.**
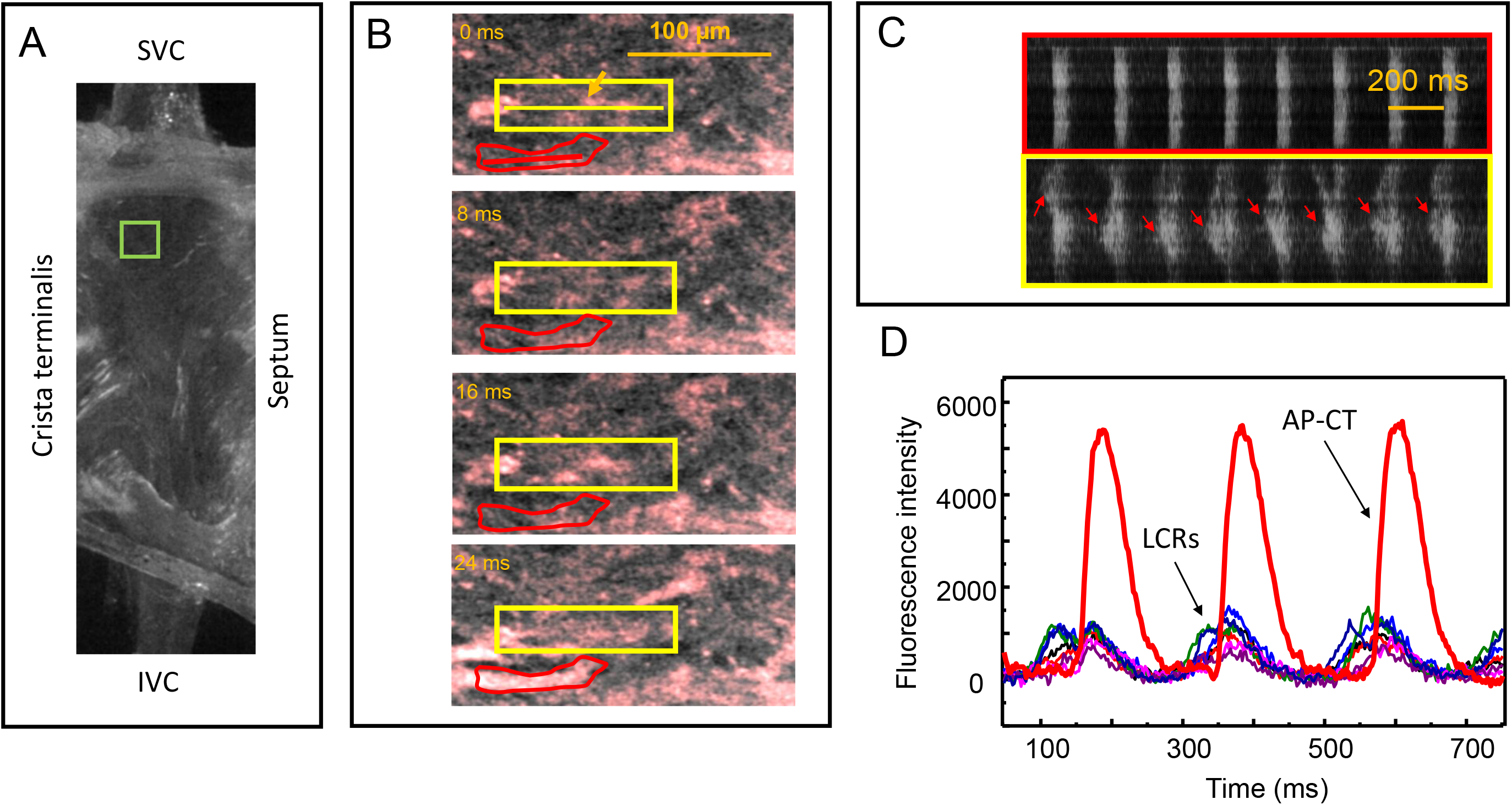
The temporal relationship between spontaneous LCRs recorded in one cell and APCTs recorded in an adjacent cell. **Panel A:** SAN image at low optical magnification. **Panel B:** A sequence of Ca^2+^ signal images from SAN cells recorded within the green ROI in panel A via a 20x water immersion objective. The cell within the yellow ROI was devoid of APCTs. Bright spots within the yellow ROI are spontaneous LCRs initiated during the diastolic phase. The cell within the red ROI was devoid of LCRs, and generated only APCTs at 16 ms. **Panel C:** Synchronized line scan images recorded along the yellow and red lines (panel B, top sub-pannel) during 15 sequential cycles within two cells highlighted by red and yellow ROIs in panel B. The colors of the frames correspond to the colors of the lines. Red arrows point to spontaneous LCRs during the diastolic phase. **Panel D:** APCTs (red color) and LCRs (traces with variable colors and smaller amplitudes) in the cell in panels B and C, plotted as function of time.

### Variable cycle-to-cycle temporal relationships of Ca^2+^ signals occurring within some SAN cells

We observed yet another type of heterogeneity of LCRs and APCTs within and among cells within the central SAN: LCRs generated in cells within the green rectangle in Fig 7A that did not fire APCTs (Fig 7B, yellow ROI, Video 6) preceded APCTs in an adjacent cell that generated both APCTs and LCRs.

**Fig 7.**
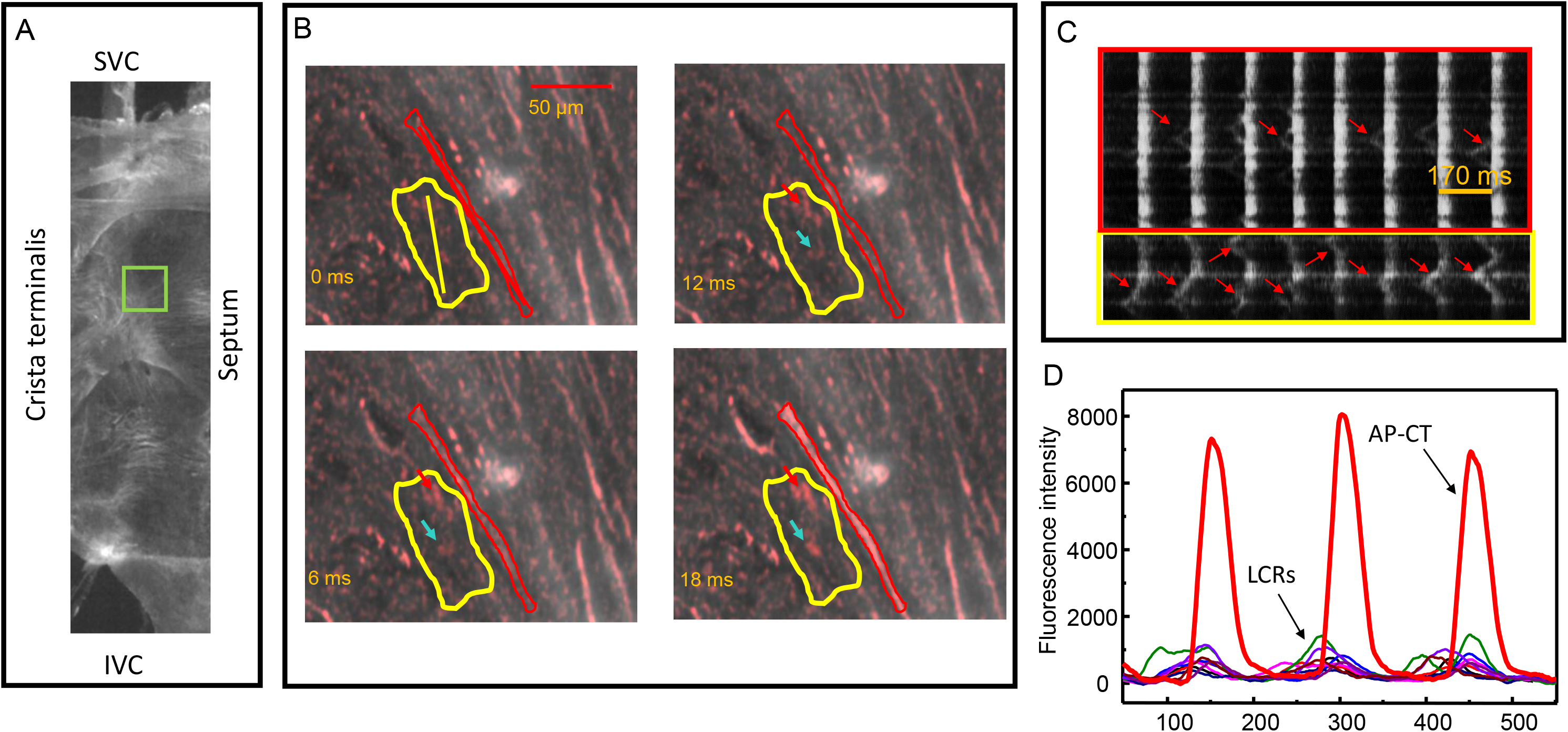
**Panel A:** Image of a SAN recorded at a low optical magnification. **Panel B:** A sequence of SAN cell Ca^2+^ signal images within the green rectangle in panel A recorded via a 10x water immersion objective. Bright spots within the yellow ROI (Panel B, red arrows) are spontaneous LCRs generated during the diastolic phase. This cell was devoid of APCTs; the cell within the red ROI generated both APCTs and spontaneous LCRs during the same AP cycles. **Panel C:** Line scan images from the two cells in panel B. The yellow and red colors of the frames correspond to the colors of ROIs in panel B. Red arrows point to spontaneous LCRs during the diastolic phase. **Panel D:** APCTs (red color) and LCRs (traces with variable colors and smaller amplitudes) in the cell with LCRs and APCTs in panel C plotted as function of time.

LCRs within the yellow ROI (red dots at 6, 12, and 18 ms) were generated during the diastolic phase of the adjacent cell within the red ROI that generated APCT at 18 ms. Line-scan images (Fig 7C) created for the two cells (Fig 7B yellow and red ROIs) confirmed that the LCRs in the cell generating no APCTs (yellow ROI) occurred prior to APCTs in the adjacent cell (red ROI). Note also that this pattern occurred during every diastolic phase. Superimposed time series of the LCRs from the cell with LCRs only (Fig 7D, thin lines with multiple colors), and APCTs from the adjacent cell having both LCR’s and APCTs (bold red line) also showed that the onset of majority of LCRs in the cell having only LCRs occurred during the diastolic phases (i.e. “entrainment zones” (14)) of the adjacent cell firing both APCTs and LCRs.

### HCN4 immunolabeling

A panorama of tiled images of a whole mount SAN preparation from superior to inferior vena cava obtained with a 2.5x objective (Fig 8) revealed the entire HCN4 expression pattern (red color) within the intact SAN, extending along the crista terminalis within the central SAN from the SVC and the septum toward the aperture of the inferior vena cava. HCN4 immunoreactivity was much stronger within the SAN than in the right atrium (to the left of the crista terminalis), in line with previous reports (28). HCN4 immunolabeling covered more than 80% of the surface of the central SAN between superior and inferior vena cava (Fig 8). Some HCN4^+^ cell clusters extended medially towards the septum and HCN4^+^ cells were also observed close to IVC, but none were detected within the crista terminalis.

**Fig 8.**
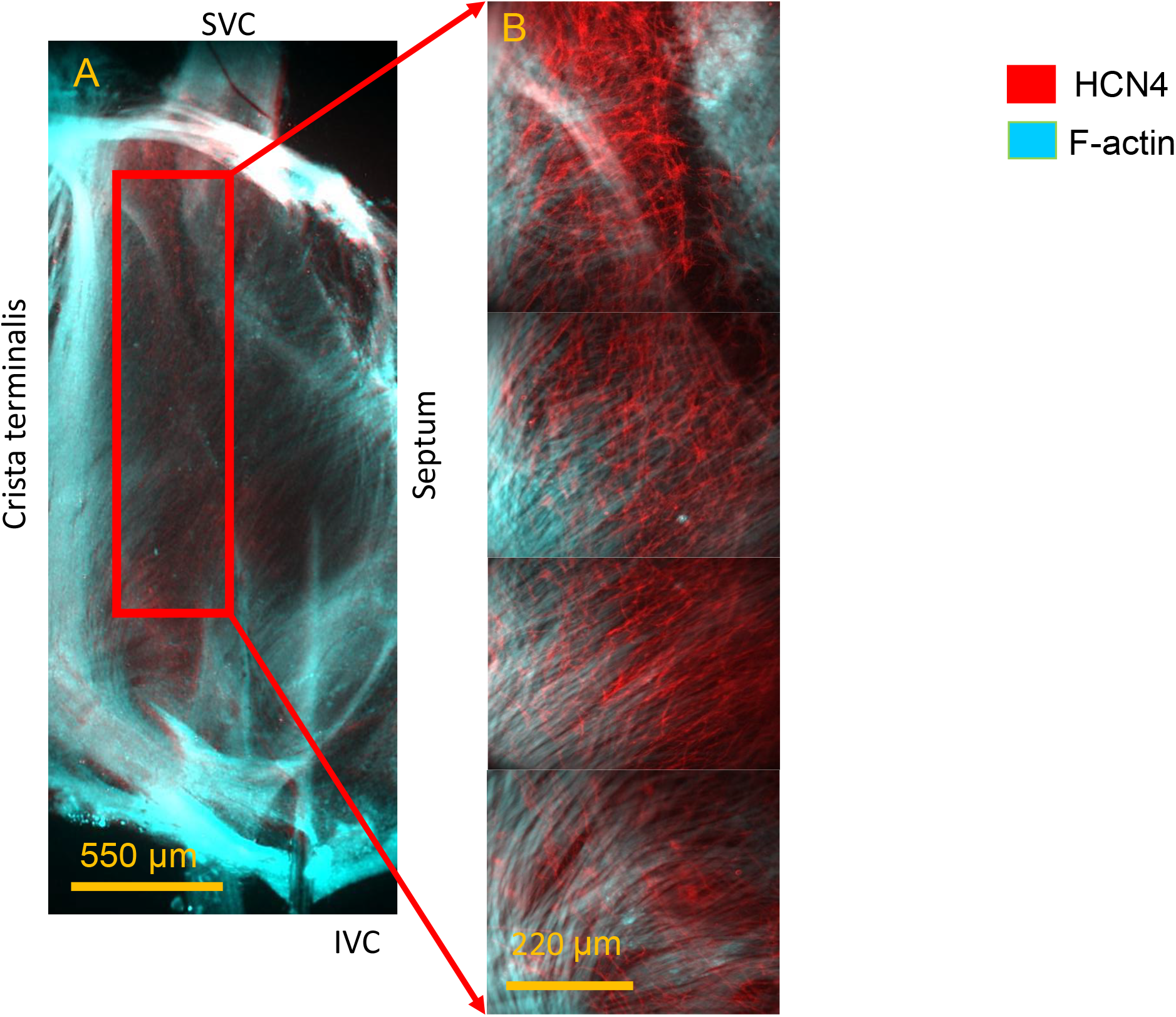
**Panel A:** An Image of a whole mount SAN preparation at low (2.5x) optical magnification, demonstrating the distribution of HCN4 (red color) immunoreactive and F-actin (cyan color) labelled cells. The merged images between HCN4 and F-actin is shown in both panels. **Panel B:** Tiled Image of the HCN4^+^/F-actin^-^ cell meshwork (red) intertwined with the HCN4^-^/F-actin^+^ cell network (cyan) reconstructed from 4 images recorded via 10x water immersion objective within the red box in panel A.

Typical HCN4^+^ cells in the central SAN visualized at higher magnification had shapes similar to those in which cell Ca^2+^ was imaged using Ca^2+^ indicator Fluo-4AM (Fig 9). Three types of cell shapes, elongated, spindle and spider type, reported previously were observed: Elongated cells had relatively uniform thickness of 2-5 μm from end to end and were 100-200 μm in length. Spindle cells were shorter and thicker than elongated with 7-15 μm thick center thinned to edges. Spider type cells had thick cell bodies of 7-15 μm and lengths of 50-80 μm and remarkably projected **multiple thin branches** towards neighboring cells (resembling to telopodes). We also identified a fourth type of cell, not described previously, having pyramidal-like cell bodies with a 15-20 μm base from which thin processes, similar to those of spider type cells, projected towards neighboring cells (Fig 9G). Thirteen cells spindle cells, 16 spider cells, 9 elongated cells and 4 pyramidal-like cells were identified within a visual field of 220μm by 220μm imaged with a 40x objective from seven optical slices taken from three SAN preparations.

**Fig 9.**
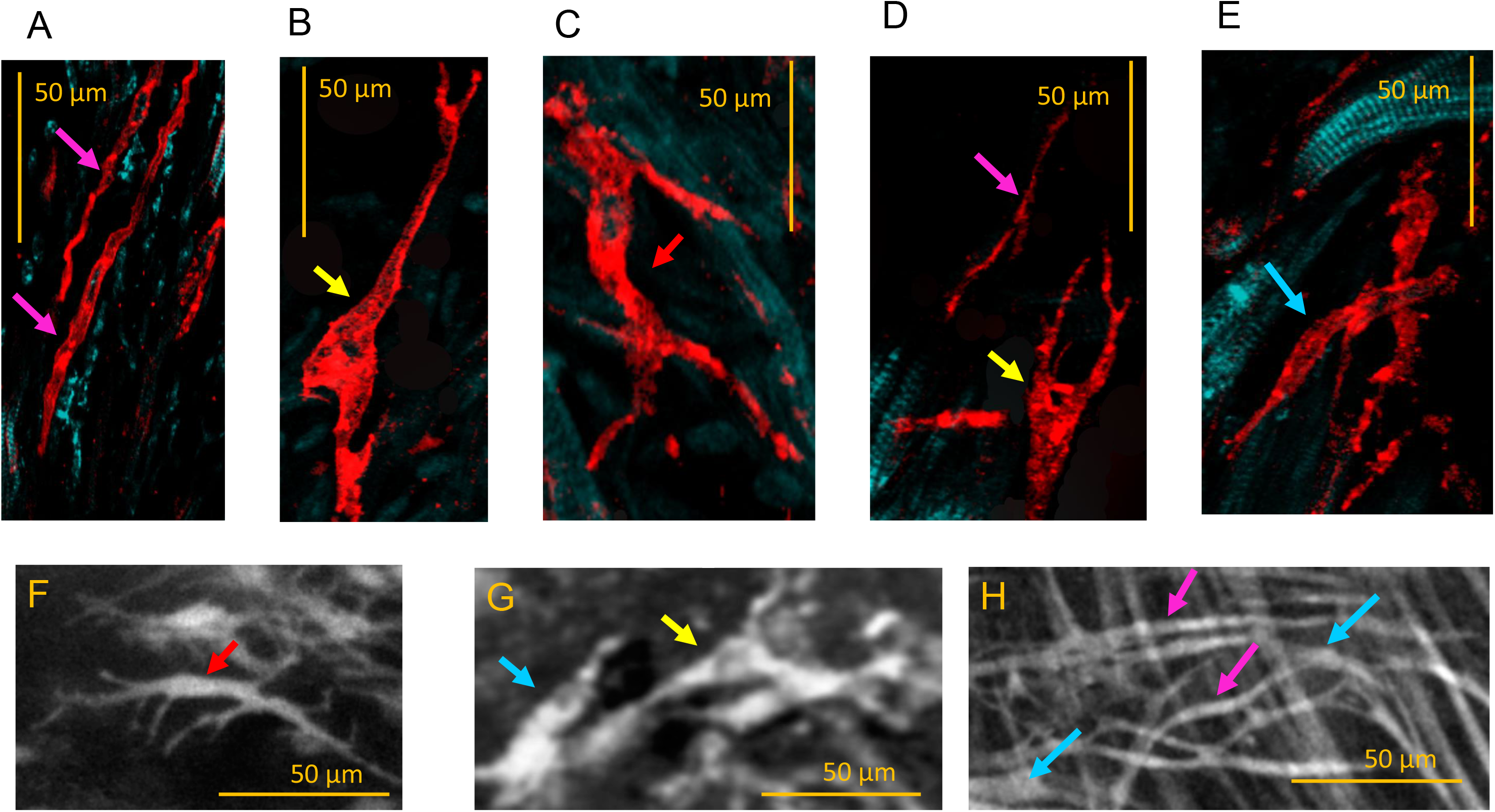
Upper panels - HCN4 immunoreactive SAN cells: elongated (magenta arrows in **Panel A**), novel, pyramidal-like shape cells (yellow arrow in **Panels B and D**), spider-like (**Panel C**), and spindle cells (**Panel E**, blue arrow). Lower panels - SAN cells loaded with Fluo-4 AM have similar shapes to immunolabelled HCN4^+^ cells in the upper panel. Spider-like cells (**Panel F**) are indicated by the red arrow. Novel cells with a pyramidal-like soma (**Panel G**) are indicated by yellow arrows. Spindle cells are indicated by the blue arrow (**Panel G and H**). Elongated cells (**Panel H**) are indicated by magenta arrows.

The cytoarchitecture of intercellular connections between HCN4^+^ cells resembled a mesh type alignment (29) in which cells within a web have numerous fine cellular branches, creating a high-density mesh of interlacing branches. Surface membranes of neighboring HCN4^+^ cells within the dense part of the continuous HCN4 meshwork were so close to each other that it was often difficult to discern cells. HCN4 labeled both the cell body and peripheral branches of these cells resembling a neurite-like arborization. Confocal imaging showed that HCN4^+^ cells were equally distributed between the epicardium and endocardium (Fig 10A-C), indicating that the HCN4^+^ cell meshwork cytoarchitecture permeated the entire thickness of the SAN.

**Fig 10.**
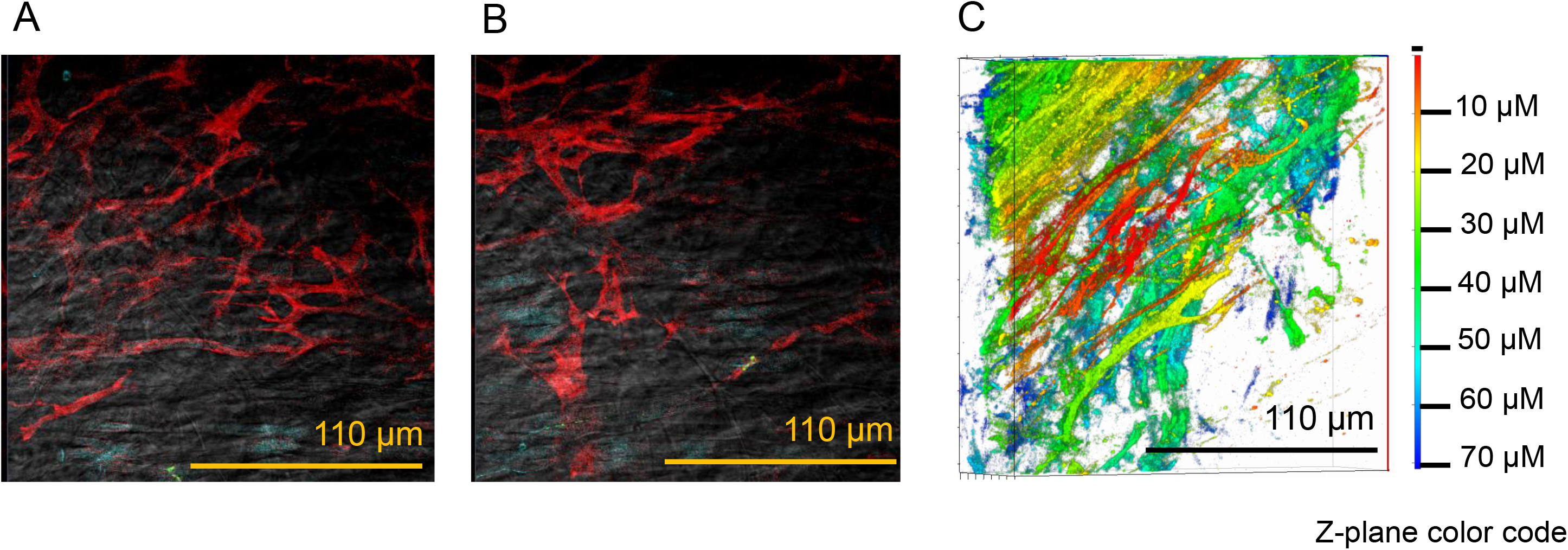
**Panels A and B**: Optical slices (1 μm thick) of HCN4 immunoreactive cells at a depth of 30 μm and 60 μm from the endocardial surface. **Panel C**: a 3D reconstruction of 70 stacked HCN4 images (optically sliced via confocal microscope) of an immunolabeled whole-mount SAN preparation demonstrating the distribution of HCN4 immunoreactive cells within a depth of 70 μM from the endocardium. Tissue depth from the endocardial site is color coded on the right side of the panel.

### HCN4 immunolabelling and co-labelling with F-actin (F-actin is not immunolabelled because we did not use antibodies)

Panoramic views of a SAN at low optical magnification (Fig 11A) of co-labeled whole-mount SAN preparations with HCN4 and F-actin (using phalloidin, a specific F-actin marker), uncovered intertwining between HCN4-expressing cells (red color meshwork) and F-actin containing cells (elongated cells in blue color network in Fig 11 B at high magnification). Although some SAN cells co-labeled positively for both HCN4 and phalloidin, the majority of HCN4^+^ cells were F-actin-negative. Note that under our imaging conditions, due to a very few F-actin filaments within HCN4^+^ cells, F-actin labeling with phalloidin was close to the background fluorescence compared to high F-actin concentration in neighboring SAN cells. Many cells, however, were F-actin-positive and HCN4-negative, clearly exhibiting two distinctly labeled networks within SAN tissue: HCN4^+^/F-actin^-^ and HCN4^-^/F-actin^+^ (Fig 11 B).

**Fig 11.**
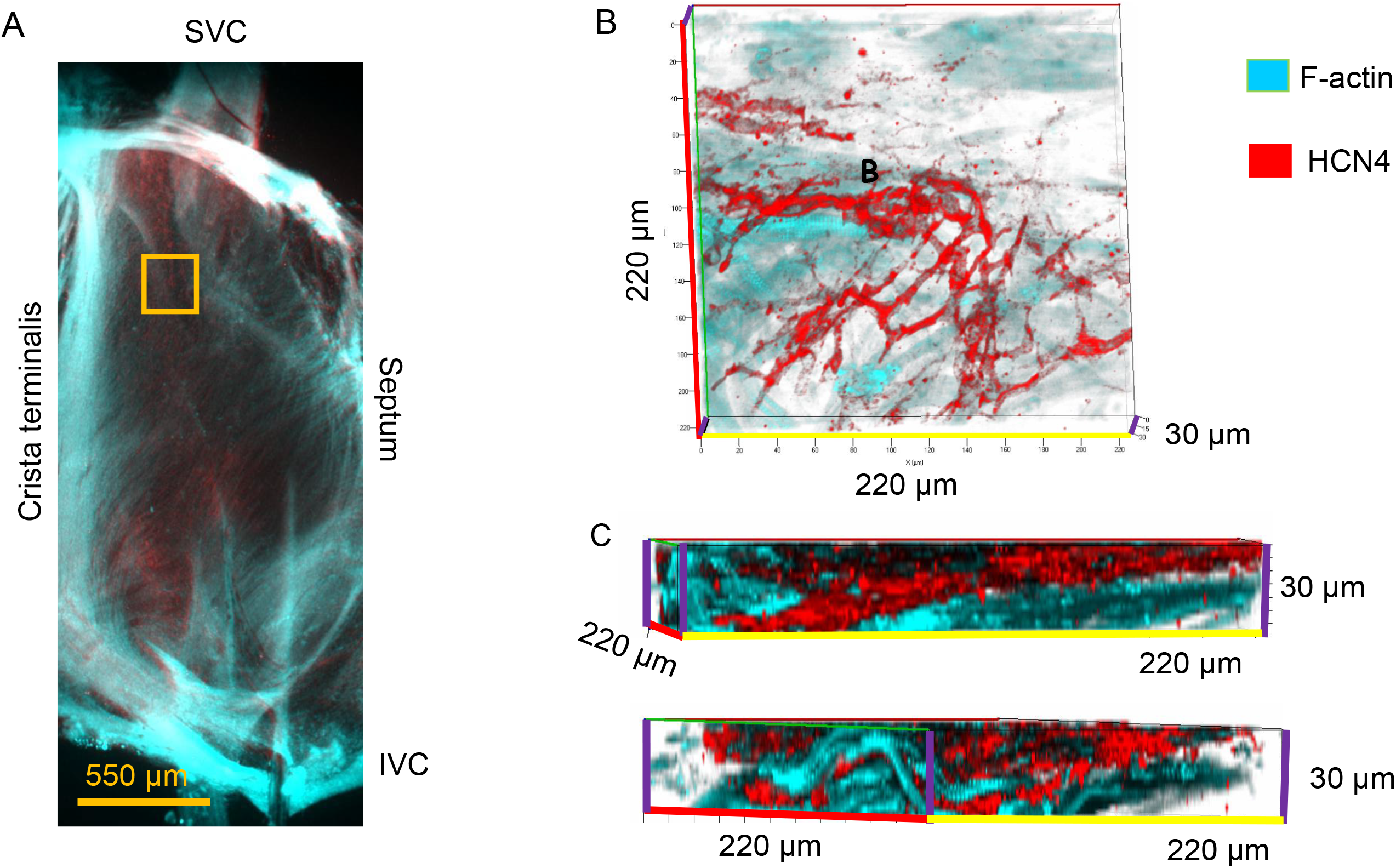
**Panel A**: An immunolabelled, whole mount image of a SAN preparation at low (2.5x) optical magnification. **Panel B:** Stacked confocal images and reconstructed front view of optically sliced z-stack images of HCN4^+^/F-actin^-^ cells (red) and HCN4^-^/F-actin^+^ cells (cyan) at a depth of 30 μm from endocardium. Confocal images were acquired with a 40x oil immersion objective within ROI (yellow) shown in panel A. **Panel C:** Side views of the z-stack images in panel B, illustrating the intertwining cells of HCN4-meshwork and F-actin networks across the 30 μm depth.

Confocal optical slicing of SAN tissue to create a series of images over sequential increments of 1 μm in depth (Fig 11) revealed the fine structural details of the intertwining of the HCN4^+^ cell meshwork and F-actin labelled cell network. Side views of the reconstructed Z-stack images in Fig 11 (Panel C) demonstrated penetration of HCN4^-^/F-actin^+^ SAN cells (blue color) into the HCN4^+^/F-actin^-^ meshwork (red color). The apparent absence of intercellular spaces between the two types of cells suggests that surface membranes of each cell are adjacent to each other. HCN4^+^/F-actin^-^ cells also penetrated the HCN4^-^/F-actin^+^ network of cells, often aligning together with HCN4^-^/F-actin^+^ cells in a 3D orientation (Fig 11B, C).

### HCN4 and CX43 co-immunolabeling

Dual HCN4 and CX43 immunolabeling of whole mount SAN preparations (Fig 12) revealed that the central SAN HCN4^+^ cell meshwork was largely devoid of CX43 (HCN4^+^/CX43^-^), and that striated HCN4-negative cells were CX43-positive (HCN4^-^/CX43^+^). Importantly, images taken in bright field fluorescent microscope showed that the HCN4^+^ cell meshwork (Fig 12 red color) and CX43 cell network (green color) become intertwined throughout the central SAN from the SVC to IVC. CX43 antibody-labeled membrane proteins observed as green dots (Fig 12C) align at the perimeter or at the ends of cells and clearly define the cell borders, revealing points of intercellular communication between CX43 expressing SAN cells. Of note, CX43 immunolabeling was not observed on the cell membranes of HCN4^+^ cells. The proximity of cells within HCN4^+^/CX43^-^ meshwork and HCN4^-^/CX43^+^ network can be observed in optical slices of the SAN obtained from intertwining areas (middle and lower image Fig 12C) in which HCN4^+^ cells come close to CX43^+^ cells but do not overlap. The continuity of CX43^+^-coupled cell network is therefore not disrupted by HCN4^+^ cells that do not express CX43 proteins, although the two cell types come close to each other.

**Fig 12.**
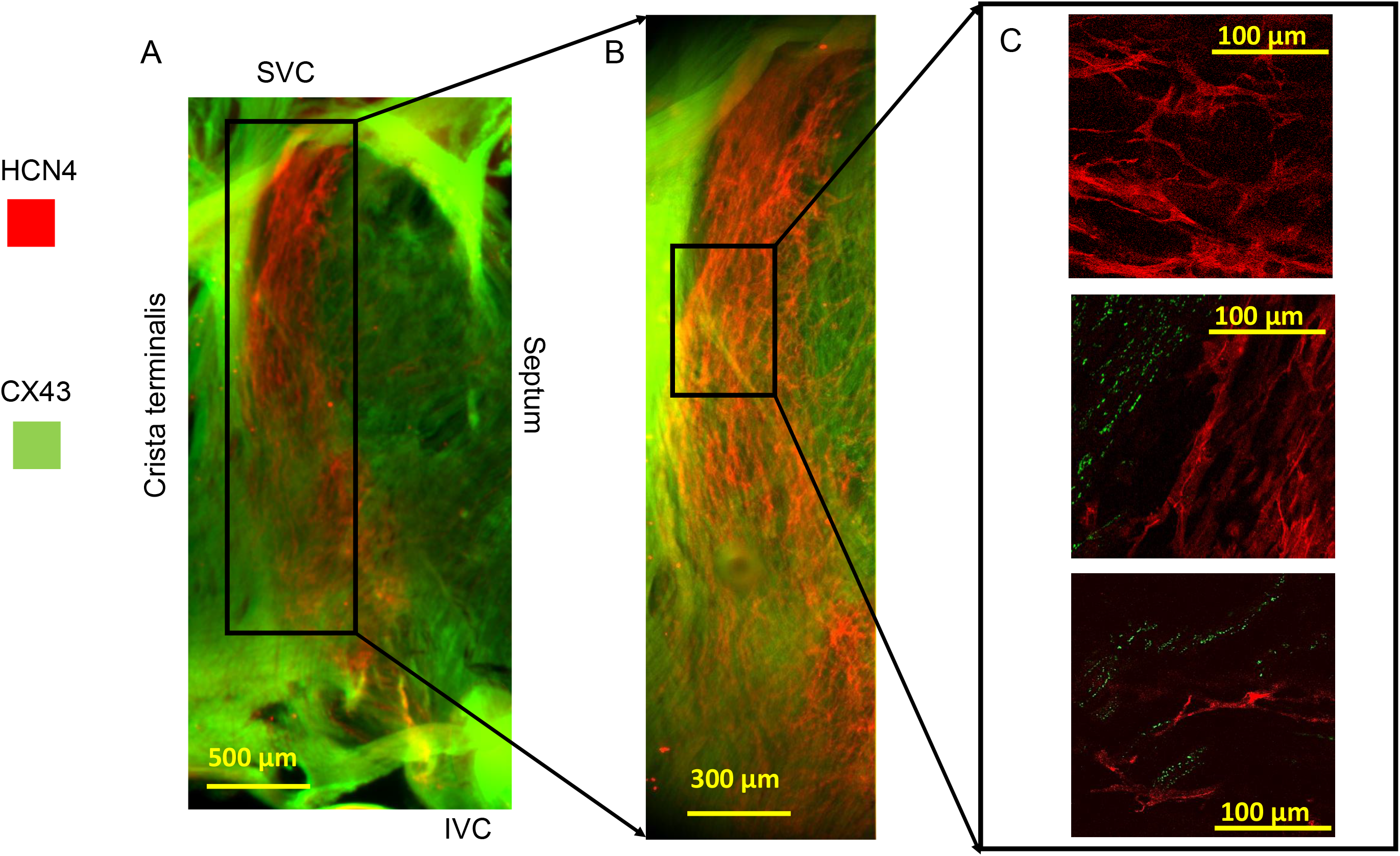
**Panel A:** A dual immunolabeled HCN4 (red) and CX43 (green) whole mount SAN image at low optical magnification (2.5x). Merged (CX43 and HCN4) immunoreactivity is shown in all three panels. **Panel B:** Image within the ROI in panel A reconstructed from 4 tile images of the HCN4^+^/CX43^-^ meshwork (red) intertwined with HCN4^-^/(CX43)^+^ network (green) taken with 10x water immersion objective. **Panels C**: Confocal images from the area within the ROI in panel B showing: HCN4^+^ cells that do not express CX43 (upper image); intertwining areas between HCN4^+^/(Cx43)^-^ meshwork (red color), and penetrating HCN4-/ CX43^+^ cells outlined by green dots corresponding to CX43 protein on the cell membranes (middle and lower panel). Note that HCN4 expressing cells in all three images do not express CX43.

### Spatial cytoarchitecture of SAN cells co-immunolabeling with HCN4, CX43 and simultaneous labelling with F-actin (Phalloidin)

Triple immunolabeling of whole mount SAN preparations with HCN4, CX43 and phalloidin antibodies revealed that intertwining between HCN4-meshwork and network of F-actin-expressing cells is not limited to specific parts of the SAN. Spatial cytoarchitecture of SAN within whole mount preparations reconstructed from 36 tiled confocal images of the area of the 1350μm by 1350μm demonstrates that areas of meshwork/network intertwining are scattered throughout the SAN from superior to inferior vena cava (Fig 13). Panoramic images showed that CX43 was expressed on F-actin containing cells but was not detected on HCN4^+^ cells. Panoramic images of large areas from intact whole mount preparations that were optically dissected into series of thin slices show that SAN cytoarchitecture cannot be described as either as mosaic or gradient: the loosely coupled HCN4^+^/F-actin^-^ meshwork is surrounded by and intertwined with an F-actin^+^/HCN4^-^ network that is strongly electrically coupled through CX43 gap junctions.

**Fig 13.**
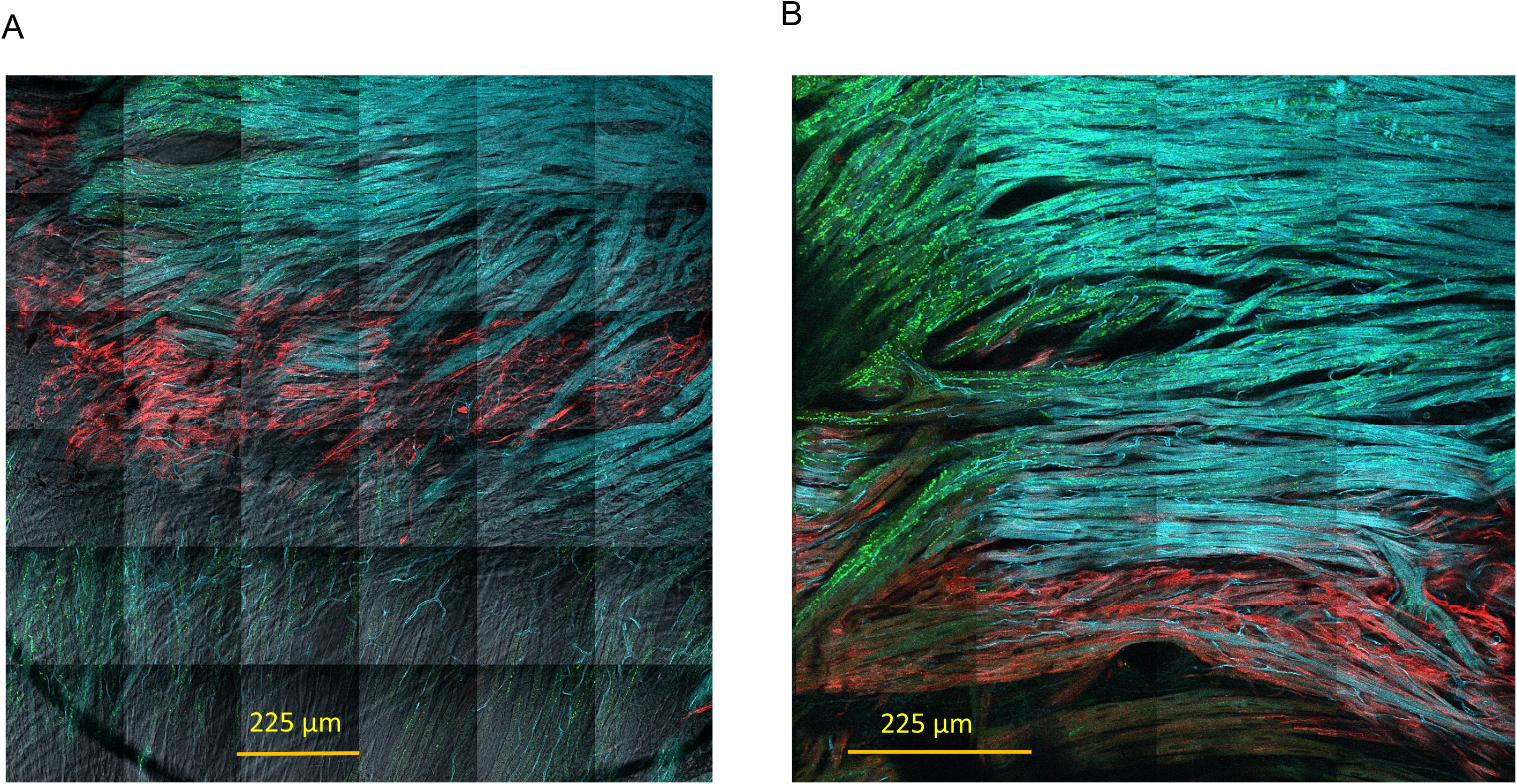
**Panel A and B:** A dual immunolabeled HCN4 (red) and CX43 (green) together with F-actin labelling (cyan) merged into a single image is shown in both panels. **Panel A:** Spatial cytoarchitecture of SAN within whole mount preparations reconstructed from 36 tiled confocal images of the area of the 1350μm by 1350μm demonstrate meshwork (red)/network(cyan) intertwining. Green dots are immunolabelled CX43 proteins. Gray color tissue was imaged in transmitted light. **Panel B:** Spatial cytoarchitecture of SAN within whole mount preparations reconstructed from 16 tiled confocal images of the area of the 900μm by 900μm of another area of meshwork (red)/network(cyan) intertwining. Green dots are immunolabelled CX43 proteins. CX43 is detected only in F-actin labeled cells.

CX43 gap junctions connect cells within F-actin^+^/HCN4^-^ network in a way that electrical signal and corresponding APCTs can propagate in directions demarcated by the CX43 protein. CX43 gap junctions appear to be randomly scattered among cells when they are colored monochromatically. However, when the position of gap junctions is color coded (Fig 14A) by depth and plotted within the z-stacks reconstructed from optical slices alignment of CX43 gap junctions making a pathway (like railroad tracks) for the transmission of electrical or Ca^2+^ signal from coupled cells is revealed. Importantly, although cells within HCN4^+^/F-actin^-^ meshwork do not express CX43 protein they may be weakly coupled through other types of gap junctions or through other non-detected intercellular coupling structures and processes that promote apparent conduction.

**Fig 14.**
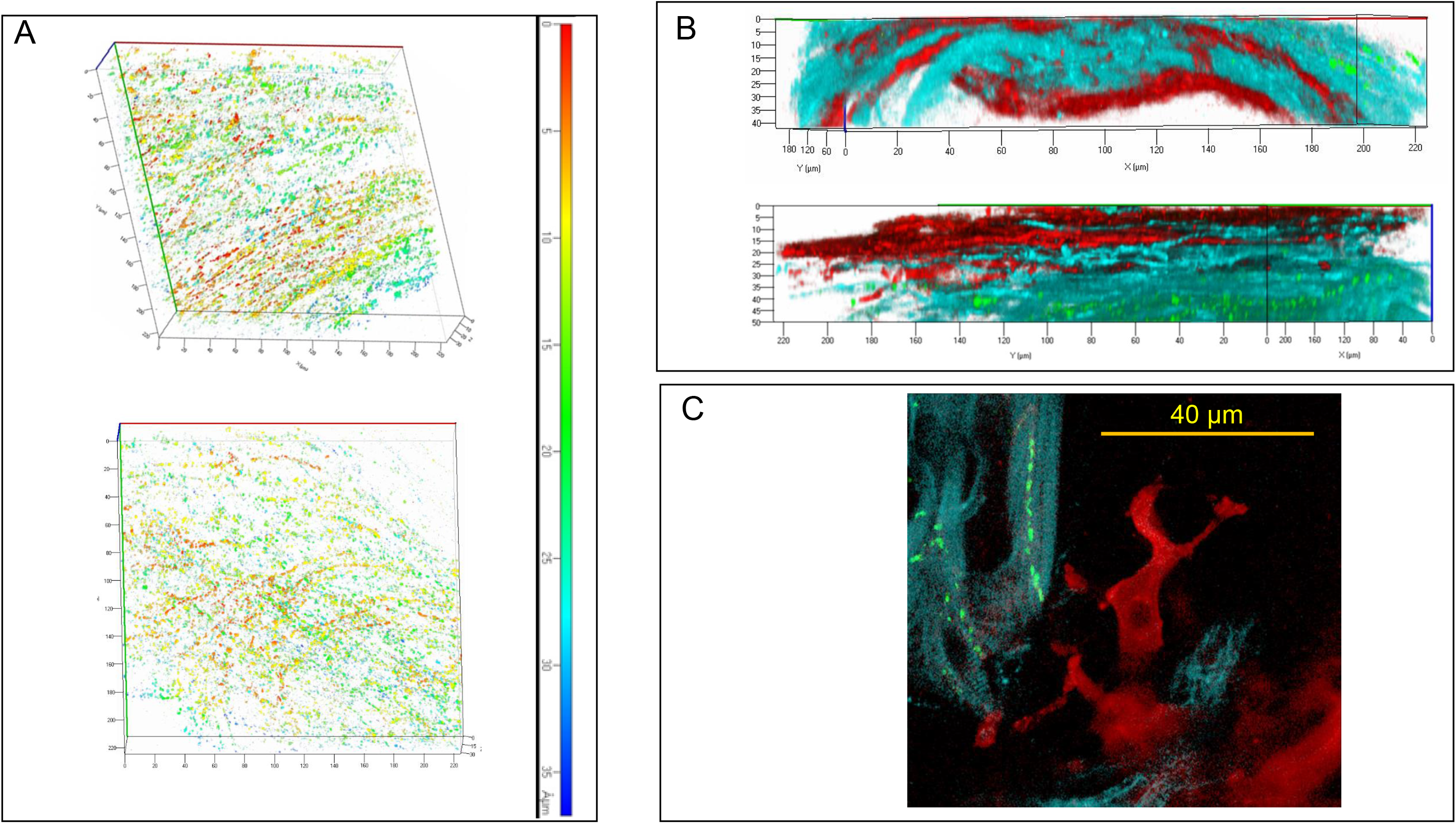
A dual immunolabeled HCN4 (red) and CX43 (green) together with F-actin labelling (cyan) merged into a single image is shown in both panels. **Panel A:** Stacked confocal images and reconstructed front view of optically sliced z-stack images of CX43 at a depth of 35 μm from endocardium. Gap junctions are color coded by depth and plotted within the z-stacks reconstructed from optical slices. Confocal images were acquired with a 40x oil immersion. **Panel B:** Stacked confocal images and reconstructed side view of optically sliced z-stack images of HCN4^+^/F-actin^-^/(CX43)^-^ cells (red) and HCN4^-^/F-actin^+^/(CX43)^+^ cells (cyan) at a depth of 40 μm and 50 μm from endocardium. Confocal images were acquired with a 40x oil immersion objective **Panel C:** Optical slice shows HCN4^+^/F-actin^-^/(CX43)^-^ cells (red) adjacent to HCN4^-^/F-actin^+^/(CX43)^+^ cells (cyan). CX43 protein (green) is expressed only in cyan cells.

### Heterogeneity of Calcium signals within SAN recorded on panoramic calcium imaging

Since the HCN4^+^/F-actin^-^ meshwork and F-actin^+^/HCN4^-^ network would be likely to have different electrical coupling capacities with respect to the presence of absence CX43, we applied a new method to represent phase heterogeneity of Ca^2+^ signaling within each pixel. Image pixels across the entire SAN were color-coded relative to the time at which the earliest APCTs occurred (i.e. chronopix). We assigned colors to each chronopix recorded at low optical magnification (2.5x objective) and defined the image as a color-coded chronopix map (Fig 15A).

**Fig 15.**
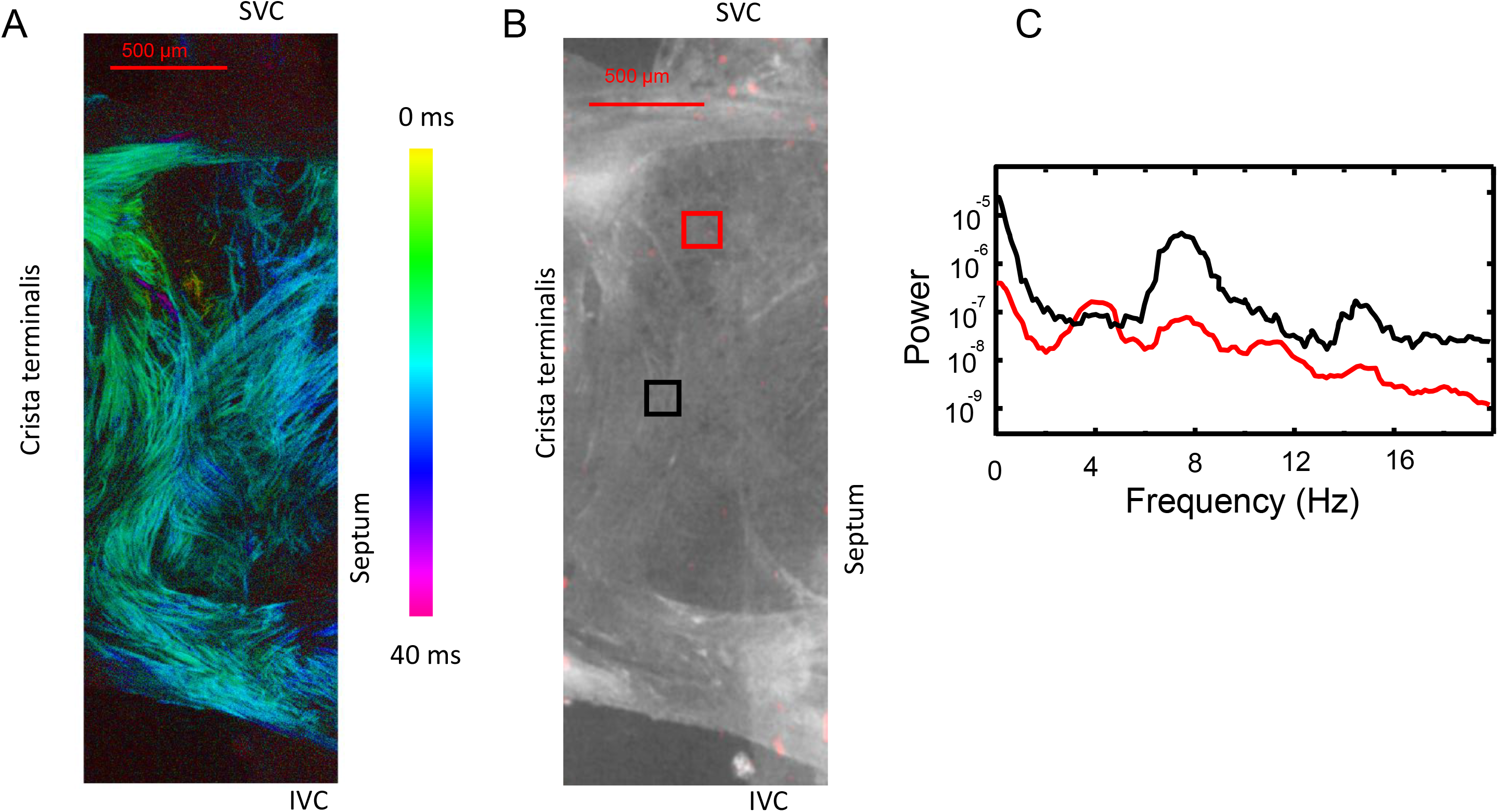
**Panel A:** chronopix of APCT emergence within the SAN color-coded in ms. **Panel Panel B:** A SAN preparation at low optical magnification: SVC-superior vena cava; IVC inferior vena cava. **C:** Superimposed plots of power spectra of Ca^2+^ signals within ROIs shown in panel B. Colors of the plots in panel C correspond to the colors of the ROI frames in panel B.

This pattern of discontinuous APCT occurrence within the panoramic chronopix image of the entire SAN demonstrated in Figure 15A is in antithesis to a concentric continuous spread observed at low resolution in previous studies (7–9) but can be interpreted as “apparent” conduction. Discontinuous APCT occurrence is due to complicated architecture of mutually intertwined HCN4^+^/F-actin^-^ meshwork and F-actin^+^/HCN4^-^ network. The HCN4^+^/F-actin^-^ meshwork manifests apparent propagation that is consistent with the idea of weakly entrained oscillators operating out of phase (12,13); while the F-actin^+^/HCN4^-^ network harbors **both** apparent and true conductional propagation.

We applied Fast Fourier transform (FFT) (Fig 15C) to detect frequency heterogeneity of low amplitude Ca^2+^ signals in pixels within HCN4^+^/CX43^-^ meshwork near to where the earliest APCTs were detected. While the power spectrum of the signal within the red ROI in panel B revealed peaks at both 4 Hz and 7.3 Hz, the power spectrum of the signal within black ROI, close to crista terminalis revealed a peak at 7.3 Hz, but a peak at 4Hz was not present indicating heterogeneous frequencies of Ca^2+^ signals within the two ROIs. It is important to note that at this low optical magnification while Ca imaging and FFT analysis resolve APCT signals and peaks of cumulative LCR activity, individual LCRs cannot be clearly detected (i.e. they are close to the noise level).

What appears to be true conductional propagation within the SAN can be observed in areas that densely stain for F-actin^+^/HCN4^-^ network (Fig 8), i.e. F-actin^+^/HCN4^-^CX43^+^ (Fig 13). We defined regions of interest (ROI) within a 2D SAN image, in which we quantified Ca^2+^ signals. Analyses of time series of APCTs recorded within different ROIs permitted assessment of heterogeneity of times of APCT occurrences among cells in which the earliest APCT and subsequent APCTs were generated (Fig 16). Ca^2+^ signal fluorescence intensity in each ROI was normalized to signal amplitude ranging from baseline fluorescence and the APCT maximum peak amplitude. The earliest detectable APCTs emerged within an area close to SVC (red circle Fig 16A). Subsequent APCTs were recorded along the crista terminalis (red, blue and orange circles) close to the point at which APCT’s occurred in the right atrium. These APCTs appeared with a delay of 9.7±2.3 ms (n= 9) with respect to the time of the earliest AP occurrence. APCTs near the septum (red, violet and cyan circles in Fig 16) appeared with a delay of 16.1+2.7 ms (n=9), respectively.

**Fig 16.**
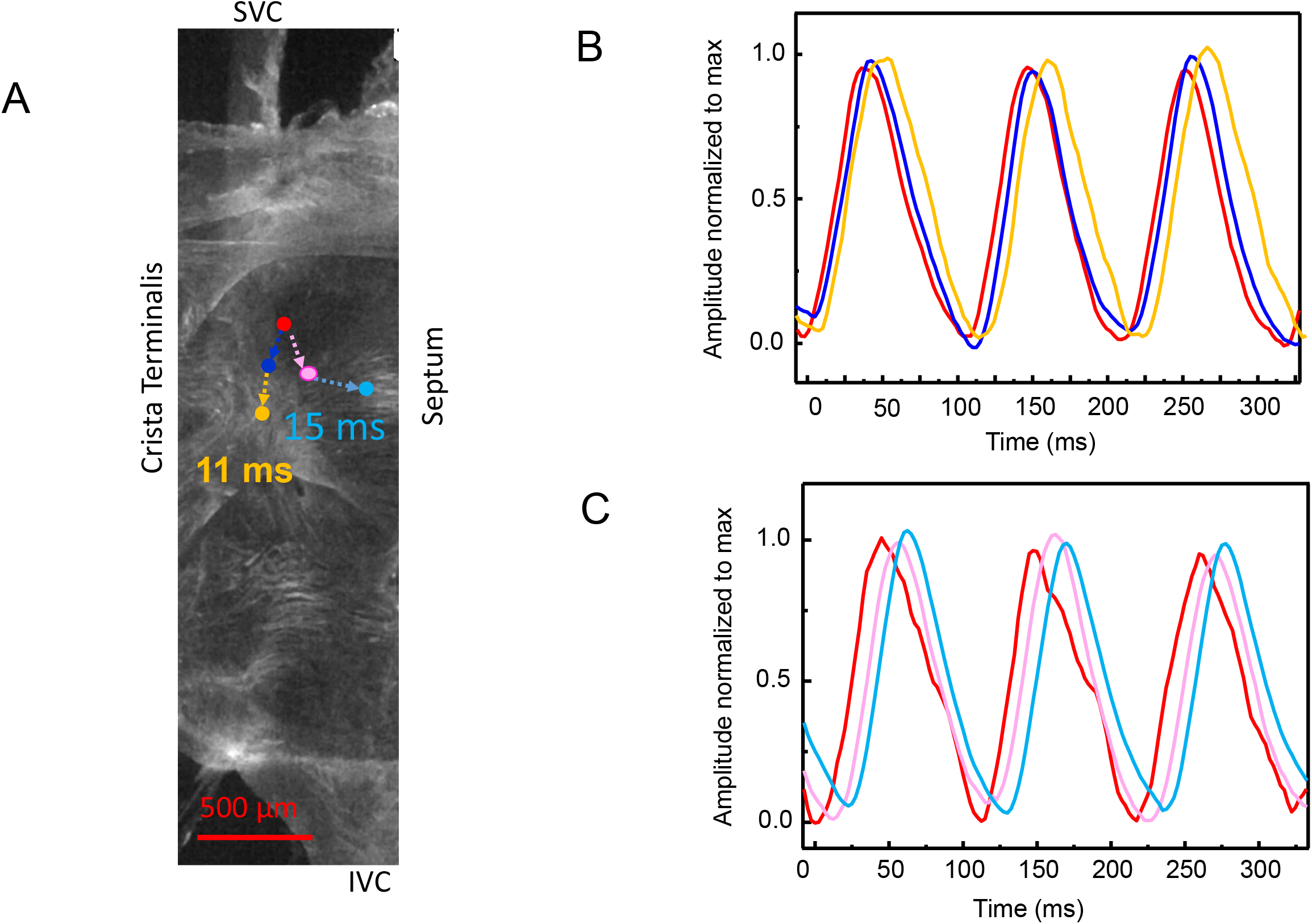
Assessment of heterogeneity of times of APCT occurrences among cells in which the earliest APCT and subsequent APCTs were generated. **Panel A:** A SAN image; SVC-superior vena cava; IVC-inferior vena cava. The dots indicate times of APCT occurrences (Numbers)(orange and light blue color ROI) following the occurrence of the earliest detectable APCT (red color ROI). **Panels B and C:** Superimposed APCTs normalized to maximum amplitude (1) over baseline fluorescence (0) and plotted as function of time. The color codes of traces correspond to the color codes of ROIs shown in panel A.

## Discussion

### Principal findings

Our results add additional insight into the structural and functional complexity of SAN tissue (1,30) (see CENTRAL ILLUSTRATION).

1. APCTs appear discontinuously within SAN with variable delays following their initial appearance close to the SVC-crista terminalis intersection.
2. A dense meshwork of HCN4^+^ cells in the central intercaval region differs in cytoarchitecture from a striated F-actin network. The HCN4^+^/CX43^-^ and F-actin^+^/CX43^+^ cells intertwine within a narrow interface zones to create **an anatomical unit** in which the intertwining areas appear to be the **functional interfaces** at which electrical and chemical signals are transmitted between the two cellular networks. This functional interface extends nearly the entire length of the SAN from the SVC to the IVC.
3. Some areas within the SAN manifest true electrical conduction while others have only apparent conduction. The HCN4^+^/F-actin^-^ meshwork manifests apparent propagation that is consistent with the idea of weakly entrained oscillators operating out of phase (12,13); while the F-actin^+^/HCN4^-^ network harbors **both** apparent and true conductional propagation. In the intertwining areas both real and apparent conduction likely to occur.
4. Oscillatory Ca^2+^ signals, including both those occurring locally, i.e. spontaneous LCRs, and those induced by APs i.e. APCTs that occur within clusters of cells comprising the HCN4 meshwork are markedly heterogeneous. In some resident SAN cells, LCRs generated in the diastolic phase appear to ignite APCTs in SAN cells, suggesting that a coupled-oscillator mechanism found in isolated pacemaker cells also operates in some cells embedded within SAN tissue. The LCR-APCT coupling among cells embedded in intact SAN tissue, however, is more complex than observed in single isolated SAN cells.
5. CX43 gap junctions connect cells within F-actin^+^/HCN4^-^ network in a way that electrical signal and corresponding APCTs can propagate in directions demarcated by the CX43 protein. When the position of gap junctions is color coded (Fig. 14A) by depth and plotted within the z-stacks reconstructed from optical slices alignment of CX43 gap junctions making a pathway (like railroad tracks) for the transmission of electrical or Ca^2+^ signal from coupled cells is revealed.

### SAN cytoarchitecture

Dual immunolabeling of whole mount SAN preparations indicated that in addition to a HCN4^+^/CX43^-^ **meshwork**, there is a **network** of F-actin^+^/CX43^+^ that intertwines with HCN4 meshwork in relatively narrow interface zones. The cytoarchitecture of HCN4 meshwork and F-actin^+^ expressing cell network however could not be categorically defined as a gradient or mosaic network (1). In contrast to HCN4 expressing cells, the striated F-actin expressing cells do not form a meshwork, but rather a network type of cells connected via Cx43, that is characterized by a repetitive pattern equal across entire central SAN. Striated CX43 expressing cells do not appear to be inserted between HCN4^+^ cells and thus do not interrupt continuity of HCN4^+^ cells within meshwork, and vice versa. Three-dimensional reconstruction of optically sliced intertwining areas in whole mount SAN preparations showed that cells from the HCN4+/F-actin-meshwork and HCN4^-^/F-actin^+^ network appear to integrate into a single structural unit. In other words, F-actin^+^ and HCN4^+^ cells were located so close to each other that an intracellular space between them could not be resolved, even with a 40x objective.

A distinction between cells within the intertwining HCN4 meshwork and F-actin network in the present study is that the majority of cells within the HCN4 meshwork are not striated and are CX43 negative, in line with the results of prior studies describing properties of the central SAN cells (26,31).

In addition to detecting three shapes: elongated, spindle and spider type HCN4^+^ cells described previously (32), novel finding of the present study is the identification of SAN cells having a pyramidal like cell body from which thin branches extended to neighboring cells within HCN4 meshwork (see Fig 9).

### Ca^2+^ signals within SAN tissue

We devised a specific experimental approach to record Ca^2+^ signals to query whether pacemaker cells imbedded in SAN tissue operate via an LCR-linked AP firing paradigm discovered in isolated cells, but on a higher, more complex, functional scale within the multi-cellular ensemble. We quantified Ca^2+^ signals across the SAN at both low and higher magnification to study fine details of LCRs with respect to AP generation manifest as APCTs.

### Discontinuous APCT occurrence across the SAN

Our first novel finding was that following earliest appearing APCTs near the SVC, APCTs appeared discontinuously in other parts of the SAN (c.f. Figs 3 and 15). This pattern of discontinuous APCT occurrence at remote cites following their earliest appearance is consistent with the idea of self-organized synchronization of loosely coupled network (33) of cell oscillators within different parts of SAN tissue (aka pacemaker cell mutual entrainment suggested by Jalife (12)). This idea of self-organized synchronization within the SAN pacemaker cell network differs from interpretations of many previous studies i.e. that pacemaker cells within an initiation site generate an impulse that spreads in a “concentric” pattern, overdriving other SAN cells that oscillate at slower spontaneous frequencies (6–9).

### Ca^2+^ signals within cells embedded within HCN4 meshwork are highly heterogeneous in spatial distribution, amplitude, frequency, and phase

Our results are, in part, consistent with the theory of mutual entrainment of SAN cell oscillators via a democratic process (12,13), but that theory considered that all cells oscillate at the same frequency but out of phase and have the same amplitude; whether subthreshold subcellular oscillations differ in amplitudes was not addressed. In other terms, the mutual entrainment theory postulated that all signals must be all or none (i.e. a full-scale AP) while firing at a common rate with the differences in phase. Thus, a second novel finding of our study is that visualization of HCN4^+^ cells in the central SAN tissue at higher magnification revealed Ca^2+^ signals that were highly heterogeneous among cells not only in phase, but also in frequency and amplitude (c.f. Fig 4). Importantly, local Ca^2+^ dynamics of cells within the HCN4 meshwork differed within clusters of cells: some cells generated only LCRs and did not fire APCTs; some only generated APCTs, manifested as synchronized bright flashes throughout the cell; and in some cells LCRs appeared during the diastolic phase prior to an APCT occurrence in that cell (c.f. Fig 5) resembling those generated by a coupled-oscillator system described in single SAN cells in isolation (19). Importantly, cells can generate APCTs both steadily and in bursts (Fig 4).

A novel type of LCR and APCT interaction was also revealed in cells within HCN4^+^/CX43^-^ meshwork, in which APCTs within a cluster of cells appear, with a delay following LCRs occurrence in **adjacent** cells that generated only LCRs, but not APCTs (see c.f. Figs 6 and 7). This spectrum of cell behaviors observed within the central SAN suggests that some cells within clusters coordinate their activity via intercellular entrainment, and different clusters of cells via inter-cluster entrainment. This crucial difference between the democratic theory of mutual entrainment (13) and our results is that we have demonstrated that cell Ca^2+^ oscillations not only differ in phase but also in frequency and amplitude. This pattern of Ca^2+^ signals differing both in phase, frequency, and amplitude among clusters of cells within SAN tissue adds a new dimension of complexity to factors involved AP generation within SAN cell networks.

Recent numerical studies of SAN network suggest importance of Ca^2+^ clock activation for robust impulse generation and resistance against annihilation (34). Our results show that LCRs in pacemaker cells embedded within SAN tissue have different entrainment patterns with respect to AP generation informed on by APCTs. One pattern is consistent with that previously discovered in single SAN cells operating in isolation in which LCRs precede APCT, confirming that the coupled oscillator theory discovered in isolated single SAN cells (19) is operational in cells within the intact SAN. Here we show that the coupled oscillator theory, however, operates at a higher and much more complex level within the network of cells comprising the SAN tissue. In other terms, we observed greater freedom and diversity with respect to Ca^2+^ cycling in cells embedded in SAN tissue than in single isolated SAN cells: many cells within SAN tissue did not fire APCTs, but generated substantial number of LCRs that varied in amplitude, frequency, and rhythmicity (see next section). While our data do not provide definitive prove that LCRs in one cell can cause events in neighboring cells, we do have a good evidence for this, i.e. a strong correlation of events in time. Indeed, in all numerous recorded cycles, LCRs in one cell preceded APCTs in the other cell. It is hardly a coincidence and points to (but does not prove) the existence of cause-effect relationship.

An additional discovery of the present study, that some cells within HCN4^+^/CX43^-^ meshwork do not fire APs, is in line with the idea put forth by Ophtof et al. (35), i.e. only a fraction of cells (~25%) embedded in SAN tissue participate in generating AP in any given electrical impulse that emanate from SAN node. We have also recently observed similar non-AP-firing behaviours in a population of enzymatically isolated guinea pig and human SAN cells: these cells have LCRs but do not fire APs (36,37). A large population of these dormant isolated SAN cells begin to fire spontaneous APs in response to increases in intracellular cAMP (36,38). Such dormant cell behaviour within SAN tissue can potentially contribute to pacemaker function via (i) having a role in entrainment of oscillators that are heterogeneous both in phase and amplitude to self-organize into synchronized signals that underlie rhythmic impulse that emanate from the SAN; and (ii) their recruitment to fire APs in response to adrenergic receptor activation or AP silencing in response to cholinergic receptor stimulation.

Immunolabeled cells embedded within the HCN4 meshwork and cells loaded with Fluo-4 within the central SAN had similar shapes. It is noteworthy that regardless their soma shape, all manifested LCRs. The topology of the HCN4 meshwork delineated by immunolabeling may be a requirement for crescendo-like self-organization of heterogeneous local Ca^2+^ signals observed in our study. In computer research meshwork properties are categorized by physical and logical topologies (39,40): physical topology relates to how the various components are placed within a network, e.g. ring, bus, mesh, or star networks; while logical topology illustrates how data flows among network components. Within a meshwork the infrastructure nodes (or infrastructure devices) connect directly, non-hierarchically to as many other nodes as possible, and cooperate with one another to route signals to the meshwork outputs. In this context, the soma of HCN4 expressing cells appear as nodes within a network with cytoarchitectural features connected via multiple branches, resembling neuronal arborization (41) that are variable and show similar topology throughout central SAN. Obscure cell borders between HCN4^+^ cells likely create contiguous pathways from cell to cell within the HCN4 meshwork. This structure is, in fact, similar to interstitial telocyte networks recently discovered in numerous tissues, including heart (42). Of note, the original name for telocytes, in fact, was interstitial cells of Cajal.

Self-organization of local signals among cells in a crescendo-like manner generates an electrical impulse that exits the SAN. This self-organization of local Ca^2+^ signals among cells within SAN tissue recapitulates the self-organization of local Ca^2+^ signals that leads to AP firing within individual pacemaker cells studied in isolation (19). This self-similar organization of local Ca^2+^ signals across scales from cells to tissue can be envisioned as a fractal-like behaviour. In other terms, while each LCR is a small, subthreshold signal, the emergent LCR ensemble signals critically contribute to generation of spontaneous APs within and among SAN cells.

### Similarities of microscale signalling within SAN and that within other tissues

The fine details of the novel micro-scale signaling paradigm within SAN tissue discovered here is reminiscent of complex information processing among clusters of neurons that create spatiotemporal synchronization of signals that drive neuronal network functions. Examples of this is type of behavior have been observed within the autonomic neural-visceral axis via its modulation of rhythms of peripheral organ function.

A characteristic of HCN4 channels, anomalous rectification, keeps diastolic membrane potential positive to the potassium equilibrium potential. Anomalous rectification is crucial for the functions of coupled oscillator systems not only for SAN cells in isolation (17,43) and in pacemaker cells embedded in SAN tissue, but also for similar systems operative in brain (44). Cells that exhibit coupled oscillator behavior within the central SAN, like many brain neurons, express HCN4.

One example occurs within the brain stem: one type of neurons, cells of preBötzinger Complex (preBötC), create spontaneous signals that activate adjacent cells (e.g. Bötzinger cells) to generate impulses that travel within nerve from the brain to the diaphragm to regulate automatic breathing (45). It has been hypothesized that only a small fraction of preBötC cells within the network is required to initiate each excitatory cycle and that the initiating preBötC cells differ from cycle to cycle (46). A major premise of this hypothesis is that spontaneous activity is initiated within a few preBötC cells and induces activity in other preBötC cells and this excitation percolates throughout the network building to a crescendo that initiates inspiration signals. We may envision that networks of pacemaker cells within SAN tissue generate local Ca^2+^ signals that are similar to those of preBötC in conjunction with signals from the heart’s little brain, i.e. the network of autonomic ganglia embedded within the atrial epicardium (47). Another example is the network of interstitial cells of Cajal that generate spontaneous Ca^2+^ signals to activate adjacent smooth muscle cells to effect gut motility (48). Our discovery of heterogeneous Ca^2+^ signals in cells comprising cardiac SAN are also reminiscent of those observed in studies in which Ca^2+^ was imaged in uterine smooth muscle (49).

### Future discoveries in SAN structure/function

Further intense study is required to understand how HCN4^+^ cells devoid CX43 signal to each other within HCN4 meshwork and how these cells signal to CX43^+^/F-actin^+^ expressing cells. The lack of structural continuity defined by cell to cell connections via one type, CX43, of gap junctions suggests a degree of autonomy of the HCN4^+^ cells from electrical behavior throughout the central SAN. Moreover, a discontinuity of CX43 expression suggests that complex Ca^2+^ dynamics within some HCN4^+^ cells may not be externally reset by AP occurrence in neighboring cells. The failure of APCTs to **reappear** within the area near the initiation site during a given AP cycle may be explained on the basis of lack of CX43 in the HCN4^+^ SAN cells that prompted the initiation of the impulse.

Whether another type of connexin besides CX43, may connect HCN4^+^ cells is a moot issue awaiting specific antibodies to other connexins; e.g. Cx45 and Cx30.2 thought to be present within the center of the SAN(26). Even if cells within HCN4 meshwork are connected via unidentified type of gap junction proteins, lack of structural continuity between CX43-negative meshwork to CX43-positive network would remain. The close proximity of two different cells types creates an interface of possible electrical or chemical communication between two networks (see Figs 11 and 14). Transmission of Ca^2+^ and electrical signals from HCN4+-meshwork to F-actin+-network would be expected to occur in these intertwining areas.

Transition of the signal between intertwined cells from HCN4^+^/F-actin^-^ cells to HCN4^-^ /F-actin^+^ cells that express different types of connexins would be expected to occur with a loss of signal amplitude and duration as reported for electrical synapses https://www.sciencedirect.com/topics/neuroscience/electrical-synapse. Indeed, sharp transitions in AP waveforms over small distances of 100 μm that occur within the mouse central SAN (50,51) are similar to differences in shapes of APs that occur during the transfer of electrical signal from pre to postsynaptic neuron through the electrical synapses (52).

Although the field of SAN biology has been dominated by the idea that electrotonic or electro-phasic impulses mediate communication among SAN cells, there is a plethora of evidence that other types of signaling occur between cells. One is mechanical (53). Other types of cell-to-cell communications, though, not yet demonstrated in SAN tissue include: ephaptic (54); photons emitted from intracellular chromophores (55,56); cellular vibrations transmitted or reflected as electromagnetic waves (57); a gas, e.g. NO, HS or CO (58); a lipid signal generated acutely by arachidonic acid degradation; as in retrograde-grade endogenous cannabinoids signaling in the brain (59); and the secretion of Ca buffers into interstitial spaces between cells (60–64). Because any of these types of cell-to-cell communications may occur among cells within SAN tissue, the foremost frontier for the field of cardiac pacemaker biology is to understand how HCN4^+^ cells communicate with each other within the HCN4 meshwork, and how they communicate signals to the CX43 cells.

**In summary**, our results demonstrate that signals generated by the SAN cell oscillators are more heterogeneous than previously formulated, including differences in amplitudes, frequencies, and discontinuities. They indicate the need for modification of the previously described elegant operational paradigm depicting resident SAN cells as a democracy involving mutual entrainment that effects a common single rate of full-amplitude APs with different local phases. Our results demonstrate novel complexity of signals of cells embedded within SAN tissue at a lower (deeper) level of events including both subcellular and whole-cell origin (i.e. within and among SAN cells), resembling neuronal networks. Synchronized macroscopic signals within the SAN, including full-scale APs, emerge from heterogeneous microscopic subthreshold Ca^2+^ signals. This signal complexity underlies robust impulse generated by the SAN and appears as a democracy at the higher level of synchronized AP generation described at a lower resolution in prior electrophysiological studies or imaging of Ca^2+^ and electrical signals.

## Acknowledgements

This work was supported by the Intramural Research Program of the NIH, National Institute on Aging. The authors thank Tracy Oppel and Loretta Lakatta for editorial assistance.

## Abbreviations list

AP: action potential
APCT: AP-induced Ca^2+^ transient
Chronopix: chrono-pixel
CX43: Connexin 43 (Gap junction alpha-1 protein)
FFT: Fast Fourier Transform
HCN4: Hyperpolarization-Activated Cyclic Nucleotide-Gated Channel 4
I_CaL_: L-type Ca^2+^ current
I_CaT_: T-type Ca^2+^ current
IVC: Inferior Vena Cava
LCR: Local Ca^2+^ Release
NCX: Na^+^/Ca^2+^ exchanger
PBS: Phosphate-Buffered Saline
preBötC: preBötzinger Complex
ROI: Region of Interest
SAN: Sinoatrial Node
SVC: Superior Vena Cava

## Video captions

**Video 1**: A low-magnification panorama of a SAN preparation from superior vena cava, SVC (aperture on the left side) to the inferior vena cava, IVC (the aperture on the right side). The distance from the SVC to IVC is about 2.5 mm. The crista terminalis is at the bottom of the videoframe. The inter-atrial septum is not visualized in the movie but is located near the top of the videoframe. Bright flashes are action potential-induced Ca^2+^ transients. As determined by phase analysis, the first APCTs occur at the red area at 2 ms in text Fig 3B.

**Video 2**: A network of cells within the SAN panorama in Movie 1 viewed at higher magnification. The length of the area captured in the videoframe is about 900 μm. Bright rhythmical flashes are action potential-induced Ca^2+^ transients that inform on the occurrences of action potentials. In addition to the bright flashes, a variety of local Ca^2+^ signals that are heterogeneous in frequency, amplitude and phase are observed (text Figure 4B, red ROIs in text Fig 4C).

**Video 3**: Heterogeneous local Ca^2+^ dynamics in the central area of SAN recorded in preparation from a genetically modified mouse pCAGGS-GCaMP8 with HCN4-targeted expression of Ca^2+^ probe.

**Video 4:** Diastolic local Ca^2+^ releases occur prior to the action potential-induced Ca^2+^ transients in the same cell over several cycles in (see text Figure 5). The length of the SAN cell in which the Ca^2+^ signals were recorded at the bottom of videoframe is about 60 μm. Local Ca^2+^ releases and action potential induced Ca^2+^ transients observed in the video were recorded within the green ROIs depicted in the text Fig 5A.

**Video 5**: A novel type of Ca^2+^ signaling in which some cells operating within a cluster (c.f. text Fig 6A green ROI) do not generate action potential-induced Ca^2+^ transients, but generate only local Ca^2+^ releases (c.f. text Fig 6B yellow ROI) that preceded action potential-induced Ca^2+^ transient firing in one or several **adjacent** cells that did not manifest their own local Ca^2+^ releases (c.f. text Fig 6B red ROI and superimposed plots in text Fig 6D). The length of the SAN cell at the bottom of the videoframe is about 85 μm.

**Video 6**: Action potential-induced Ca^2+^ transients (APCTs) occur out of phase in two cell clusters (upper right and middle of video) recorded from the green ROI in the central SAN depicted in text Fig 7A. Note, in the cell cluster in the middle of the video the self-organization of local Ca^2+^ releases prior to APCTs in the adjacent cells within the cluster. Note also that local Ca^2+^ releases also appear within the cell that generates APCTs, but prior to APCT firing.

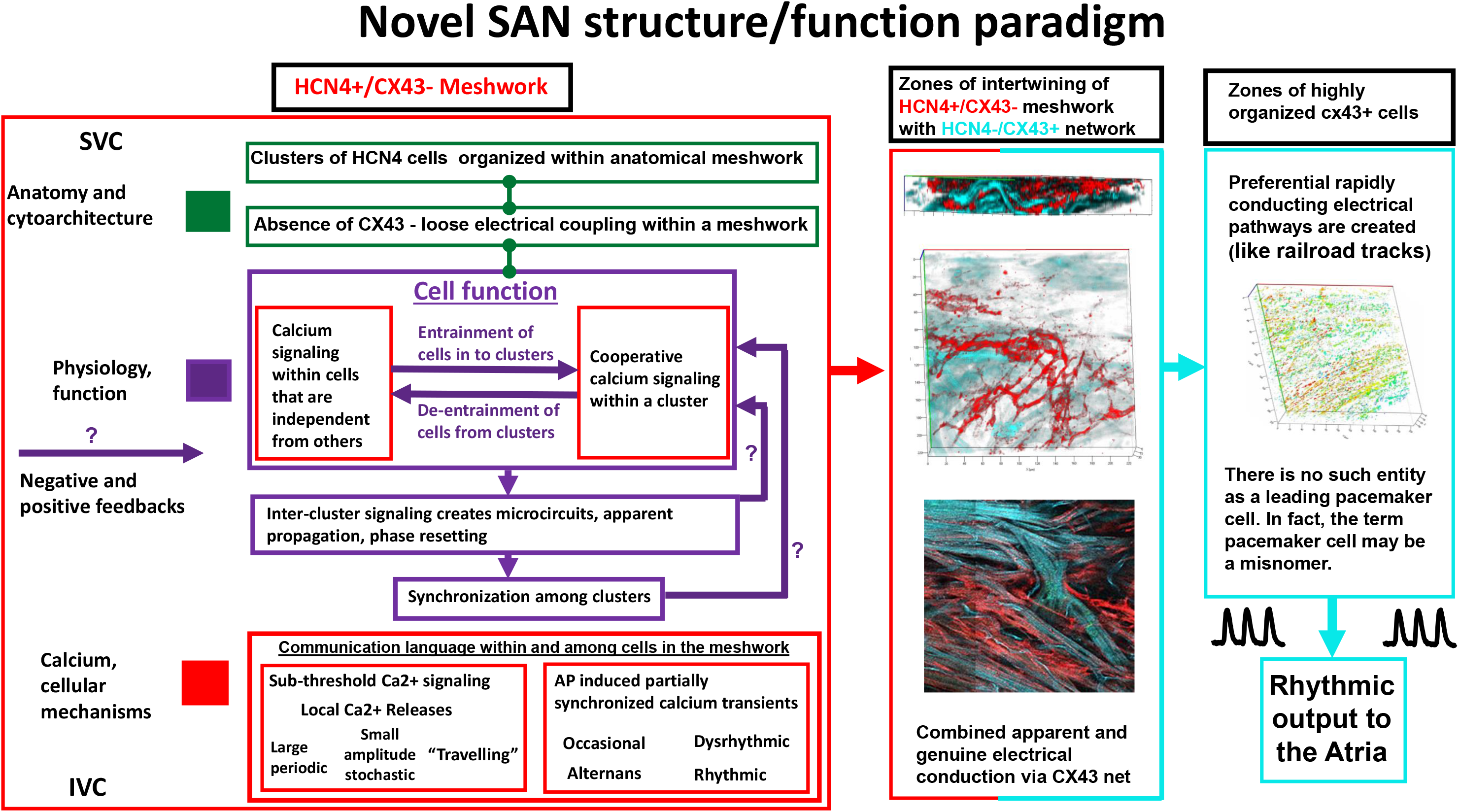

